# Robust CRISPR Screens Identify TPL1 as a Novel Long Noncoding RNA Driving Triple-Negative Breast Cancer Hallmarks

**DOI:** 10.1101/2025.08.06.668327

**Authors:** Ramesh Elango, Sunandini Ramnarayanan, Radhakrishnan Vishnubalaji, Lutfiye Yildiz Ozer, Michela Coan, Yifan Yu, Sunkyu Choi, Tina Uroda, Carlos Pulido-Quetglas, Frank Schmidt, Khalid Ouararhni, Ayman Al Haj Zen, Rory Johnson, Nehad M. Alajez

**Affiliations:** Translational Oncology Research Center (TORC), Qatar Biomedical Research Institute (QBRI), Hamad Bin Khalifa University (HBKU), Qatar Foundation (QF), PO Box 34110, Doha, Qatar; School of Biology and Environmental Science, University College Dublin, Dublin, D04 V1W8, Ireland; Conway Institute for Biomolecular and Biomedical Research, University College Dublin, Dublin, D04 V1W8, Ireland; The SFI Centre for Research Training in Genomics Data Science, Dublin, Ireland; College of Health & Life Sciences, Hamad Bin Khalifa University (HBKU), Qatar Foundation (QF), Doha, Qatar; Proteomics Core, Research, Weill Cornell Medicine-Qatar, Education City, Qatar Foundation, P.O. 24144 Doha, Qatar; Department of Medical Oncology, Inselspital, Bern University Hospital, University of Bern, Bern 3010 Switzerland; Department for BioMedical Research, University of Bern, Bern 3008, Switzerland.; Graduate School of Cellular and Biomedical Sciences, University of Bern, Bern 3012, Switzerland; Genomics Core Facility, Hamad Bin Khalifa University, Qatar Foundation, Doha P.O. Box 34110, Qatar; FutureNeuro, SFI Research Centre for Chronic and Rare Neurological Diseases, Dublin, Ireland

**Keywords:** Long noncoding RNA (lncRNA), Triple-negative breast cancer, CRISPR-Cas9 screen, TPL1, ceRNA network

## Abstract

Despite the growing catalog of long noncoding RNAs (lncRNAs), the functional roles of their vast majority in cancer remain poorly defined. To systematically explore lncRNA dependencies in triple-negative breast cancer (TNBC), we compiled a comprehensive annotation by merging GENCODE, BIGTranscriptome, and MiTranscriptome databases and performed a CRISPR-Cas9 deletion screen targeting 1,029 TNBC-enriched lncRNAs. The screen revealed several essential lncRNAs and those modulating doxorubicin sensitivity, with TPL1 emerging among top hits. TPL1 silencing significantly impaired TNBC cell proliferation in both 2D and 3D cultures and reduced invasive capacity in an organ-on-chip model. Transcriptomic and proteomic profiling following TPL1 knockdown revealed downregulation of genes involved in ECM–receptor interaction, focal adhesion, cell migration, and PI3K-Akt signaling. Mechanistically, TPL1 directly interacted with key proteins including EIF4B, MDM2, TARBP2, TLE5, and GTPase RAN, suggesting TPL1 could regulate RNA processing, transcriptional repression, and translation, as well as modulate GTPase signaling pathways. Additionally, TPL1 functioned as a competing endogenous RNA (ceRNA), sequestering miR-10396b-5p, miR-486-3p, and miR-450a-2-3p, among others, thereby modulating expression of pro-tumorigenic targets. Clinically, TPL1 was significantly overexpressed in TNBC tissues, particularly in the BLIS subtype. Collectively, our findings highlight TPL1 as a key regulator of TNBC molecular networks and a promising therapeutic target.

## Introduction

Triple-negative breast cancer (TNBC), accounting for 15-20% of all breast cancers, lacks estrogen receptors (ER), progesterone receptors (PR), and human epidermal growth factor receptor 2 (HER2), making it unresponsive to hormonal or anti-HER2 therapies. Therefore, chemotherapy remains the primary treatment, with Anthracyclines and Taxanes being the most commonly used agents. Novel therapies like Poly (ADP-ribose) polymerase (PARP) inhibitors, including olaparib and talazoparib, are approved for BRCA-mutated TNBC ^1,2^. Immune checkpoint inhibitors, such as pembrolizumab and atezolizumab, have also shown promise ^3^. However, a significant proportion of patients still experience relapse, highlighting the need for better understanding of the disease and development of new therapeutic approaches ^4^.

Long non-coding RNAs (lncRNAs) have emerged over the past decades as key regulatory elements under health and disease settings ^5^. Unlike their protein-coding counterparts, lncRNAs, characterized by lengths exceeding 200 nucleotides, do not encode for functional proteins; rather, they exert their influence through diverse mechanisms in regulating transcription, chromatin modification, and post-transcriptional processing ^6–8^. Altered expression of lncRNAs has been linked to various human diseases, including cancer ^9^. According to GENCODE release (R) 47, approximately 35,900 lncRNA genes have been annotated. Although the number of lncRNAs is likely to be much higher, due to the increasing efforts of comprehensive annotation and sequencing of the non-coding part of the transcriptome ^7^. While many lncRNAs have been reported thus far, the functionality of only a slight proportion of annotated lncRNAs has been described and has been linked to various human diseases ^10^. CRISPR-Cas9 genomic screens have revolutionized the search for novel therapeutic targets in cancer, through systematically perturbing genes in cancer cells to identify those essential for survival, growth, or drug resistance ^11–13^. While this approach proven successful in studying protein-coding genes (PCGs), the same concept may not be applicable to lncRNA research since a single indel is unlikely to have drastic effects on those non-coding molecules. Esposito et al. recently used a CRISPR-Cas9 deletion screen targeting lncRNA transcription start sites (TSSs) to identify lncRNA-based dependencies and therapeutic vulnerabilities in lung cancer, supporting RNA-based therapeutic interventions ^14’15^.

While previous studies have primarily focused on annotated lncRNAs, our study uniquely maps the functional landscape of both annotated and novel lncRNA candidates in TNBC. Using a genome-wide CRISPR-Cas9 deletion screen targeting > 1,000 lncRNAs, we identified key regulators of TNBC cell proliferation and doxorubicin resistance. Among them, TPL1 emerged as a critical lncRNA associated with extracellular matrix organization, cellular metabolism, and GTPase activity, processes central to TNBC progression. Functional validation with antisense oligonucleotides (ASOs) confirmed that TPL1 silencing significantly impaired cell proliferation and invasion, highlighting its therapeutic relevance. Mechanistic analyses revealed that TPL1 functions both as a protein scaffold, interacting with cancer-associated proteins such as EIF4B, MDM2, and RAN and as a competing endogenous RNA (ceRNA) that sponges key microRNAs including miR-10396b-5p, miR-486-3p, and miR-450a-2-3p. Through these dual mechanisms, TPL1 orchestrates gene networks that drive TNBC aggressiveness, positioning it as a promising target for future RNA-based therapeutic strategies.

## Materials and Methods

### Library construction and cloning

To construct a comprehensive transcriptome assembly, we first retrieved Gene Transfer Format (GTF) annotations from the GENCODE R19 (GRCh37), BIGTranscriptome ^16^, and MiTranscriptome ^17^ and constructed a merged assembly using StringTie version 1.3.3b using the ‘-merge’ option ^18^. We then employed a strict strategy to select lncRNA and miRNA candidates for library construction. Firstly, we excluded candidates with TSS either overlapping a PCG transcript or within <5kb up- or downstream of any PCG TSS, as per the GENCODE v38liftv37 and the aforementioned Stringtie annotation. We also employed a second filter to remove candidate TSSs that were close to a PCG exon up or downstream (1.5 kb downstream or 3.5 kb upstream from candidate TSS) that may have escaped the first filter. Secondly, we computed transcript expressions using a panel of TNBC cell lines (MDA-MB-231, BT-549, MDA-MB-157, MDA-MB-468, HCC70, and MDA-MB-453) RNA-Seq data. Transcripts exhibiting TPM ≥ 0.1 in at least 3 cell lines were subsequently retained. FANTOM peaks were obtained from MDA-MB-453 TNBC cell line and from all other cell line data to correct the candidate TSSs obtained from the GTF files. If the TSS of a candidate was within 50 bp of a FANTOM peak, the TSS is replaced with the location of the FANTOM peak, giving priority to FANTOM peaks in TNBC cells over peaks obtained from all cell lines. Remaining TSSs that were ≤ 50 bp proximity were grouped into single TSS clusters. For neutral control pgRNAs, genomic regions with no anticipated impact on cell phenotype were designated as described before with extra checks to ensure that the neutral controls did not overlap with any gene in the stringtie GTF ^14^.

CRISPETa ^19^ was employed to generate approximately ten unique pgRNAs for each candidate TSS cluster region. During the design phase, the CRISPETa software’s off-targeting filters were applied. Finally, we were able to design 10 pgRNA for 869 targets, 9 pgRNA for 1 target, 8 pgRNA for 136 targets and less than 8 pgRNAs for 23 targets. The final library design comprised 11,695 unique pgRNA, with 9896 pgRNAs targeting 1,029 unique candidate TSSs (945 unique candidate genes including genes with merged TSSs, 894 unique genes when the merged TSSs are separated), 599 pgRNAs (97 AAVS1 + 502 pgRNAs for 51 unique intergenic or intronic regions) neutral controls, and 600 pgRNAs targeting the open reading frame (ORF) and 600 pgRNAs targeting the TSS for each of the 60 positive control genes. The positive control genes were chosen from DepMap CRISPR gene effects for TNBC cell lines. Genes with the most negative score (mean score calculated for 6 cell lines used in the study) were prioritized. Library cloning was done by VectorBuilder (https://en.vectorbuilder.com/) using Mammalian Dual-gRNA Expression Lentiviral Vector (VB160903-1018vur, Figure S1).

### Cell line authentication by STR

Genomic DNA (gDNA) extracted from the two TNBC models used for the screen (MDA-MB-231 and BT-549) was used as input for STR profiling using AmpFLSTR Identifiler PCR amplification kit (Thermo Fisher Scientific, Inc., Waltham, MA, USA) following the manufacturer’s protocol. Briefly, 1 ng of gDNA was subjected to PCR amplification. Positive and negative controls were run in parallel with the samples. After amplification, PCR products were prepared for electrophoresis by adding Hi-Di Formamide and size standard mix to each sample and allelic ladder. Electrophoresis was performed in the Genetic Analyzer 3500xl DX system. Allelic call analysis was performed on Gene Mapper software from Applied Biosystems (Thermo Fisher Scientific, Inc., Waltham, MA, USA). Reference STR profiles for the cell lines were retrieved from ATCC (https://www.atcc.org/).

### Doubling time calculation

Cas9-expressing MDA-MB-231 and BT-549 cells were seeded in triplicate at 1×10^5^ cells/well in 6-well plates. The cells were allowed to grow for 48 and 72 hr, respectively. Subsequent to this growth period, the cells were enumerated, and their doubling times were determined using an online calculator available at https://www.omnicalculator.com/biology/cell-doubling-time. The calculated doubling times for the MDA-MB-231 and BT-549 cell lines were determined to be 30.0 ± 0.4 and 25.2 ± 0.3 hr, respectively, based on a sample size of three (n = 3).

### Generating stable Cas9-expressing TNBC cell models and cell sorting

Lentivirus production of Cas9-expressing lentiviral plasmid was carried out by co-transfecting HEK293FT cells with 2.5 µg of Cas9 plasmid carrying blue fluorescent protein (BFP) and blasticidin S-resistance gene (Addgene 78545), and 2.5 µg of Delta 8.91 and 250 ng of the packaging pVSV-G plasmids as described before ^20^ using Lipofectamine 3000, according to manufacturer’s protocol. The supernatant containing viral particles was harvested at 48 and 72 hr after transfection and was used to transduce MDA-MB-231 and BT-549 TNBC models using spin centrifugation. The human MDA-MB-231 and BT-549 TNBC cells were cultured in Dulbecco’s Modified Eagle Medium (DMEM) supplemented with 10% fetal bovine serum (FBS) and 1% penicillin/streptomycin (Pen-Strep), all were purchased from Thermo Scientific (Thermo Scientific, Rockford, IL, USA). Cells were cultured as an adherent monolayer at 37 °C under 5% CO2 in a humidified incubator. Cells were then expanded, selected in blasticidin S (20 µg/ml for MDA-MB-231 and 10 µg/ml for BT-549) for 7 days, then were sorted twice to enrich for MDA-MB-231 and BT-549 highly expressing Cas9 using BD FACSAria III (BD Biosciences Inc., San Diego, CA, USA) to achieve > 85% BFP-positive cells. Flow cytometric assessment of BFP (Cas9) and GFP (pgRNA) expression was conducted using BD LSRFortessa X-20 flow cytometer (BD Biosciences, CA, USA).

### pDECKO Cloning of Candidate 256 Using Gibson Assembly

Dual-guide RNAs (gRNAs) targeting Candidate 256 were designed using the CRISPETa tool to ensure compatibility with the U6 and H1 promoters followed by cloning as described before ^21^. The sequences of the two gRNAs targeting Candidate 256 (LINC00910) are listed in Table S1. First, the pDECKO_mCherry plasmid (Addgene #78534) was linearized using BsmBI-v2 (NEB #R0739S) at 37°C for 2 hours. A 155 bp gRNA-containing oligonucleotide fragment was then assembled into the digested vector backbone using Gibson Assembly (NEB #E2611L) at 50°C for 1 hour. Two microliters of the assembly reaction were transformed into DH5α competent E. coli (Invitrogen #18258012), resulting in an intermediate plasmid harboring the gRNA pair but lacking the constant sgRNA scaffold and H1 promoter. In the second step, the intermediate plasmid was re-digested with BsmBI-v2 at the Insert-1 site to linearize the vector for insertion of the remaining components. To prevent self-ligation, the digested product was dephosphorylated and purified to eliminate residual enzymes and small DNA fragments. The constant region and H1 promoter (Insert-2), derived from pDECKO-MALAT1 (Addgene #72622), were then assembled into the purified backbone via Gibson Assembly. The final assembly products were transformed into E. coli, and resulting colonies were screened by colony PCR. Positive clones were confirmed by Sanger sequencing.

### Lentiviral titration

High titer lentiviral particles (>10^9^ TU/ml) from the pooled library were prepared by Vector Builder (VectorBuilder Inc., Chicago, IL, USA). Prior to lentiviral transduction, titration experiments were performed. Briefly, 2×10^6^ MDA-MB-231 or BT-549 TNBC cells were seeded in 12 well plate in DMEM supplemented with 8 µg/ml polybrene (Sigma-Aldrich, St. Louis, MO, U.S.A). Subsequently, lentiviral ranging from 0.25 to 4.0 µl were added to each well followed by spin-infection at 37° C, 1200 rpm for 90 min ^14^. After centrifugation, the media was replaced with fresh media without polybrene, and cells were incubated overnight. Twenty-four hr later, the cells from each well were counted and were split into two equal aliquots, of which one was treated with puromycin (2 µg/mL). After 72 hr, the viral titer was calculated by dividing the number of surviving cells in the puromycin well, by the number in the puromycin-free well.

### Lentiviral transduction and CRISPR-del proliferation and doxorubicin-resistance screen

The pooled CRISPR-del library screen was conducted as described before ^14^. Briefly, MDA-MB-231 and BT-549 cells expressing Cas9 were transduced with the pooled lentiviral particles at multiplicity of infection (MOI) of 0.5 using spin centrifugation for 90 min at 37° C in the presence of polypyrene (8µg/ml). Transduced cells were then selected for 5 days with puromycin (2 µg/ml) prior to the initiation of the screen. After selection, cells from each experimental replica were pooled and 12×10^6^ cells were collected as time point 0 (T0). Twelve million cells (∼ 1,000x coverage) were then subjected to 2-dimentional (2D) culture in the absence or presence of low dose (7.5 nM) doxorubicin. Cells were passaged every 2-3 days, while maintaining the same 1,000x coverage until end of the experiment (T2: 21 doublings).

### gDNA preparation and sequencing

gDNA was isolated from 12×10^6^ cells using the Qiagen Blood and Cell Culture DNA Midi Kit (#13343, Qiagen Inc., Hilden, Germany) following the manufacturer’s protocol. The concentrations of gDNA were determined using a NanoDrop 2000 spectrophotometer (Thermo Scientific, DE, USA). For PCR amplification, 40 µg of each sample was used. Each reaction (4 µg x 10 replicas) was set up in 100 µl reactions, comprising 10 µl (4 µg) of gDNA, 50 µl of Collibri™ 2X Library Amplification Master Mix (cat# A38539250, Invitrogen), 5 µl of forward universal primer (10 µM), 5 µl of a uniquely barcoded reverse primer (10 µM), and 30 µl of nuclease-free water. PCR cycling conditions were as follows: initial denaturation at 98°C for 30 sec, followed by 25 cycles of 15 sec at 98°C, 30 sec at 52°C, 30 sec at 72°C, with a final extension at 72°C for 5 min. The forward and reverse oligos for PCR and NGS of the library are provided in Table S1. PCR products were purified using AMPure XP beads (#A63880, Beckman Coulter, Brea, CA, USA), and the purified product was quantified using Qubit dsDNA HS Assay Kits (ThermoFisher #Q32854). Subsequently, samples underwent NGS (2×150 bp) on Illumina NextSeq™ 4000, with approximately 50 million reads acquired for each sample.

### Screen hit identification and prioritization

Amplicon sequencing reads were analyzed over several steps to ensure quality check, process, and to quantify library sequences, before differential analysis to identify drop out hits. Read qualities are tested by FastQC and outputs were stored for inspection (http://www.bioinformatics.babraham.ac.uk/projects/fastqc/). Sequencing reads (FASTQ files) were first trimmed using cutadapt ^22^ based on adapters that are either specified by the users or set by default. Trimmed reads are then mapped to the indexed protospacer library using STAR ^23^. Gene significance was assessed using MAGeCK ^24^. This was performed using the published CASPR pipeline ^25^. CASPR is designed to analyze CRISPR screen using pgRNAs, it takes four input files: fastq-format read files, library of single or paired gRNAs, experimental design, and controls. Command CASPR -f “{forward.fastq.gz” -r "{reverse}.fastq.gz” --library library.txt -y 0.25 -k -- output-dir./results --controls controlfile.txt --exper-design expdesign.txt was used to perform CASPR analysis. The FDR threshold was set to 0.25. Volcano and box plots were generated using R. The plots depict both negative (red) and positive (blue) selections from MAGeCK. The significance thresholds used were a p-value cutoff of 0.25 and a log fold change cutoff of -0.3 and 0.3. Circos plot presentation was done using SRplot, as previously described ^26^.

### Antisense oligonucleotide transfection and colony formation unit (CFU) assay

Antisense oligonucleotides (ASOs) targeting the respective lncRNA candidate were designed using Qiagen oligonucleotide designer (https://geneglobe.qiagen.com/). Five different ASOs were designed for each candidate. Cell transfection was carried out using reverse transfection approach and lipofectamine 2000 (Invitrogen) as we previously described ^27^. Briefly, 0.48 µl of ASO (30 nM final) was diluted in 50 μL of Opti-MEM (GIBCO, Carlsbad, CA, USA), and 0.6 μl of Lipofectamine 2000 was diluted in 50 μl of OPTI-MEM. The diluted ASO and Lipofectamine 2000 were mixed and were incubated at room temperature for 20 min. Two hundred µL of transfection mixture was added to the tissue culture plate (12-well plate), and subsequently 1.6 x 10^5^ MDA-MB-231 or BT-549 cells in 600 μL transfection medium (complete DMEM without Pen-Strep) were added to each well. After 24 hr, the transfection cocktail was topped up with complete DMEM. For CFU assay, transfected cells were kept in culture for 7 days, then were washed twice using phosphate-buffered saline (PBS). Subsequently, plates were stained with crystal violet (0.1% in 10% ethanol) and the plates were then placed on shaker for 2-3 hr. Plates were then air-dried at room temperature before imaging, and CFUs were quantified by dissolving crystal violet in 10% SDS, and subsequent measurement of absorbance at 590λ.

### Three-dimensional (3D) organotypic viability assay

Viability assays in 3D cultures were conducted as we previously described ^27^. Briefly, BT-549 cells transfected with ASOs against the indicated candidate lncRNAs or negative control ASO were pelleted on day 4 and were subsequently combined with Growth Factor Reduced (GFR) Basement Membrane Matrix Matrigel (Corning Corp., Bebford MA, USA) at density of 3.0 x 10^4^ cells/mL. Subsequently, several drops of the cell suspension were dispensed into pre-warmed (37 °C) 60 mm Ultra-low attachment culture plates (Corning Corp., Bebford MA, USA). The plates were then inverted and placed in a 37 °C, 5% CO2 cell culture incubator to allow the droplets to solidify for 20 min before adding 4–5 mL of growth medium and further incubation. Organoid growth formation was observed under the microscope on the indicated dates.

### Time-course proliferation assay

GFP-labeled MDA-MB-231 cells ^28^ were reverse-transfected with ASOs to assess their impact on cell proliferation. Briefly, Part A (0.12 μL of ASO in 12.5 μL of OPTI-MEM) was mixed with Part B (0.15 μL of Lipofectamine 2000 in 12.5 μL of OPTI-MEM), and the mixture was incubated at room temperature for 20 minutes to allow complex formation. Following this, 7,000 MDA-MB-231-GFP cells suspended in 75 μL of complete DMEM (lacking Penicillin/Streptomycin) were seeded into each well of a 96-well plate containing 25 μL of the transfection mixture, yielding a final ASO concentration of 30 nM. After 24 hours, the transfection medium was replaced with fresh complete DMEM, and cells were incubated for an additional 72 hours at 37°C in a humidified incubator with 5% CO₂. At 96 hours post-transfection, cells were fixed with 4% paraformaldehyde and stained with Hoechst 33342 and CellMask-IR for nuclear and cytoplasmic visualization, respectively. Imaging was performed using the Operetta High-Content Imaging System (PerkinElmer) at 10× magnification. Nine fields per well were captured, and cell numbers were quantified using Harmony image analysis software. Proliferation data were normalized to the cell count at time zero for each corresponding well.

### Scratch wound healing assay

To evaluate the effect of ASO-mediated knockdown on cell migration, GFP-labeled MDA-MB-231 cells were reverse-transfected and subjected to a scratch (wound healing) assay. The transfection procedure was identical to that described for the proliferation assay: Part A (0.12 μL of ASO in 12.5 μL of OPTI-MEM) was mixed with Part B (0.15 μL of Lipofectamine 2000 in 12.5 μL of OPTI-MEM) and incubated at room temperature for 20 minutes to allow complex formation. Following complex formation, 25,000 MDA-MB-231-GFP cells per well were seeded in 75 μL of complete DMEM (without Penicillin/Streptomycin) into wells containing 25 μL of the transfection mixture, achieving a final ASO concentration of 30 nM. After 24 hours, the media was replaced with fresh complete DMEM, and cells were incubated until they reached confluency (approximately 72 hours post-transfection). A uniform scratch was introduced into the confluent monolayer using the AutoScratch™ Wound Making Tool (BioTek Instruments). At 36 hours post-scratch, cells were fixed with 4% paraformaldehyde and stained with Hoechst 33342 for nuclear labeling and Phalloidin-568 for actin filament visualization. Imaging was performed using the Operetta High-Content Imaging System (PerkinElmer) at 10× magnification, capturing nine fields per well at 24-hour intervals. Wound (gap) area was quantified using Harmony image analysis software, and the area remaining at 36 hours was normalized to the initial wound area at time zero as described before ^28^.

### Tumor cell invasion using OrganoPlate^®^-3-lane system

To evaluate the effect of ASO-mediated knockdown on MDA-MB-231-GFP cell invasion, the OrganoPlate^®^ 3-lane platform (Mimetas) was used ^28^. A 4 mg/mL type I collagen solution was prepared by diluting a 5 mg/mL rat tail collagen I stock (R&D Systems) and neutralizing it with 10% 37 g/L Na₂CO₃ (pH 9.5) and 10% 1 M HEPES buffer. Then, 2.2 μL of the collagen I gel was loaded into the middle lane of the chip and incubated at 37°C with 5% CO₂ for 15 minutes. To promote cell adhesion, 40 μL of fibronectin (10 μg/mL in PBS) was added to the outlet of the perfusion channel and incubated overnight. GFP-labeled MDA-MB-231 cells were reverse-transfected in a T25 flask. For transfection, Part A (3.84 μL of ASO in 400 μL OPTI-MEM) was mixed with Part B (4.8 μL of Lipofectamine 2000 in 400 μL OPTI-MEM), incubated at room temperature for 20 minutes, and combined with 4.8 mL of complete DMEM (without Penicillin/Streptomycin) containing 480,000 cells, resulting in a final ASO concentration of 30 nM.

After 24 hours, cells were harvested and resuspended in complete DMEM at a density of 8,000 cells/μL. The perfusion channel was washed with PBS, and 4 μL of the cell suspension was seeded into the inlet of one perfusion channel. The plate was tilted on its side inside the cell culture incubator for 2–3 hours to promote cell attachment. Subsequently, 50 μL of DMEM was added to both the inlet and outlet of the seeding side, while 50 μL of EGM2 growth medium was added to the opposite inlet and outlet to establish a chemoattractant gradient for directional invasion. The plate was then placed on the Mimetas rocker platform in the incubator (14° angle, 8-minute interval) to initiate bidirectional flow. At 120 hours post-transfection, cells were fixed with 4% paraformaldehyde and stained with Hoechst 33342 and Phalloidin-568. Imaging was performed at 10× magnification using the Operetta High-Content Imaging System (PerkinElmer) at 24-hour intervals. For each chip, spinning disk confocal imaging was conducted with 40 z-steps (5 μm spacing). Four adjacent fields were acquired per channel to capture the entire invasion area. Image stacks were stitched and segmented using Harmony imaging analysis software. For time-course analysis, the number of invaded cells at each time point was normalized to the initial (time zero) cell count. To assess endpoint invasion, the number of invaded cells at 120 hours post-transfection was normalized to the total cell number in the tumor channel.

### Viability staining using fluorescent microscopy

Fluorescence microscopy was employed for the detection of cell death. Briefly, the acridine orange and ethidium bromide (AO/EtBr) fluorescence staining method was employed, where TNBC cells under different treatment conditions were incubated for 5 days. Subsequently, cells were washed twice using PBS before staining with a dual fluorescent solution containing 100 μg/mL AO and 100 μg/mL EtBr (Sigma Aldrich, St. Louis, MO, USA) for 2 min. The stained wells were then observed and imaged using an Olympus IX73 fluorescence microscope (Olympus, Tokyo, Japan). AO staining was used to visualize cells undergoing apoptosis, while EtBr-positive cells indicated the presence of necrotic cells.

### RNA isolation and reverse transcription-quantitative polymerase chain reaction (RT–qPCR)

RNA was extracted from transfected cells using the miRNeasy Kit (Qiagen Inc., Hilden, Germany) following the manufacturer’s instructions. The concentration and purity of the isolated RNA were assessed using NanoDrop 2000c (Thermo Scientific, MA, USA), and the RNA samples were stored at −80 °C. For reverse transcription, 1 µg of total RNA was utilized using the high-capacity cDNA reverse transcript kit (Applied Biosystems, Foster City, CA, USA). The quantification of lncRNA expression was conducted using PowerUP SYBR Green Master Mix and a QuantStudio 7 Flex real-time PCR system (Applied Biosystems, Foster City, CA, USA), following the manufacturer’s protocol. The relative fold change (FC) in lncRNA expression was determined using the 2−ΔΔCt method, where the average of ΔCt values for the target amplicon was normalized to Actin Beta (ACTB) endogenous control and compared with control samples. The primer sequences used for gene expression and lncRNA quantification and genotyping are listed in Table S2, and their design was carried out using Primer3 (https://www.ncbi.nlm.nih.gov/tools/primer-blast/).

### NGS and RNA-Seq analysis

MDA-MB-231 cells transfected with the indicated ASOs targeting specific candidate lncRNAs underwent total RNA isolation at 72 hr post-transfection. The quality and quantity of the isolated RNA were assessed using on-chip electrophoresis employing the Agilent RNA 6000 Nano Kit (Agilent Technologies, CA, USA) and the Agilent 2100 Bioanalyzer (Agilent Technologies), following the manufacturer’s guidelines. Samples with RNA Integrity Number (RIN) > 8 were considered for library preparation using the TruSeq Stranded Total RNA Library kit (Illumina Inc., San Diego, CA, USA) according to the manufacturer’s protocol. In brief, 100 ng of total RNA underwent rRNA depletion according to manufacturer’s protocol. The initial cDNA synthesis involved random hexamers and SuperScript II Reverse Transcriptase (Thermo Fisher Scientific, Waltham, MA, USA). The second cDNA strand synthesis substituted dTTP with dUTP. The resulting double-stranded cDNA was end-repaired and adenylated, followed by ligation of barcoded DNA adapters to both ends and subsequent amplification. Library quality was assessed using an Agilent 2100 Bioanalyzer system, and quantification was performed using the Qubit system. The libraries were then pooled, clustered on a cBot platform, and sequenced on an Illumina HiSeq 4000, generating approximately 50 million paired-end reads (2 × 75 bp) per sample.

To confirm candidate lncRNA depletion in ASO-treated samples, the abundance of targeted lncRNAs were estimated relative to GENCODE R45 using Kallisto 0.42.1. For mRNA transcriptome quantification, pair-end FASTQ files were subsequently aligned to the GRCh38 reference genome using the built-in module and default settings in CLC genomics workbench v21.0.5. Expression data (total counts) were imported into iDEP.951, normalized (CPM, counts per million), and were subjected to data transformation using EdgeR (log2(CPM+c)), as previously described ^27^. DESeq2 was employed to identify Differentially Expressed Genes (DEGs) in iDEP.951.

### RNA-Seq data retrieval from TNBC cohort and abundance estimation

Raw sequencing data from 360 TNBC and 88 normal tissue were retrieved from the Sequence Read Archive (SRA) database (accession# PRJNA486023) ^29^ using the SRA toolkit v2.9.2 ^30^. The Kallisto index was constructed from our merged assembly by creating a de Bruijn graph in kallisto 0.42.1 using default settings ^31^. Abundance estimation of each lncRNA was quantified using kallisto 0.42.1 and the corresponding index file as we described before ^32^. Normalized expression data in TPM (transcript per million) were then imported into iDEP.951 and transcripts with minimum expression of 0.1 TPM in at least 10 samples were retained as we described before ^27^.

### Correlation analysis between TPL1 and protein-coding genes in TNBC

To investigate the transcriptome-wide association between TPL1 (ENSG00000289985) and other protein-coding genes (PCGs), we conducted a correlation analysis using bulk RNA-seq data from a cohort of 360 triple-negative breast cancer (TNBC) samples. Gene expression values were obtained from a processed expression matrix in TPM format, where each row represented a gene and each column a patient sample. Using Python (v3.9) and the pandas and scipy libraries, we computed pairwise Pearson correlation coefficients between TPL1 expression and all other genes. Specifically, TPL1 expression values (first row of the matrix) were used as the reference, and the Pearson correlation coefficient (r) and associated p-value were calculated against each gene across the cohort using the scipy.stats.pearsonr() function. To ensure robustness, missing or undefined values were handled by substituting with default values (r = 1.0, p = 0.0) where appropriate. The final output included gene-wise correlation coefficients and p-values, which were compiled into a single DataFrame and exported as a CSV file for downstream analysis and visualization. All analyses were performed in a reproducible computational environment using Python.

### Gene set enrichment analysis (GSEA) and modeling of gene interactions

Differentially expressed genes identified from the RNA-Seq analysis were imported into Ingenuity Pathway Analysis (IPA) software (Ingenuity Systems, USA) for functional regulatory network and canonical pathways analyses. Utilizing Upstream Regulator Analysis (URA), Downstream Effects Analysis (DEA), Mechanistic Networks Analysis (MNA), and Casual Network Analysis (CNA) prediction algorithms as previously described ^27^. IPA utilizes a precise database to form functional regulatory networks from individual gene lists. Each network is assigned a statistical score, the Z score, based on its fit to the set of focus genes. Biological functions associated with each network are ranked according to their significance.

### Analysis of TPL1 lncRNA–protein interactions using TLC-CLIP data

RNA–protein interaction data were retrieved from the Targeted Ligation and Crosslinking Crosslinking and Immunoprecipitation (TLC-CLIP) dataset (PRJNA935334). FASTQ files were mapped to the GENCODE v47 reference genome using the CLC Genomics Workbench. Transcript expression levels of TPL1 (ENSG00000289985), measured in TPM, were extracted from each sample and used to compare binding by hnRNPA1, hnRNPC, and hnRNPI relative to RBFOX2. A detailed description of the methodology is available in the published study by Ernst et al. ^33^.

### Data-independent acquisition mass spectrometry (DIA-MS) proteomic analysis

MDA-MB-231 cells were transfected with either TPL1-targeting ASO1 or a non-targeting control ASO. After 72 hours, cells were lysed and proteins were extracted for analysis. Data-independent acquisition mass spectrometry (DIA-MS) was conducted using a Vanquish Neo UHPLC system coupled to an Orbitrap Exploris 480 (Thermo Scientific). Peptide samples (500 ng) were loaded via autosampler, trapped on an Acclaim PepMap Nano-Trap column, and separated on an EASY-Spray PepMap Neo C18 column over a 120-minute gradient (3–40% buffer B, 0.1% formic acid in 80% ACN) at 250 nL/min. Ionization was performed using an EASY-Spray Source at 2.2 kV and 45 °C.

Full MS scans were acquired at 120,000 resolution over a 350–1200 m/z range, followed by DIA MS/MS scans with a 35 m/z window, 30,000 resolution, and 30% HCD.

Raw data were processed using SpectroPipeR (v0.4.2) in R (v4.4.3). Protein quantification was based on normalized MS2 peak area intensities, with median normalization applied when Spectronaut™ internal normalization was not used. Zero intensity values were imputed using half the minimum intensity of the dataset, and methionine-oxidized peptides were excluded. Peptide intensities were calculated by summing ion intensities per peptide per sample. Protein-level quantification was performed using the MaxLFQ algorithm (via the iq R package), followed by global median normalization. Principal component analysis (PCA) and UMAP dimensionality reduction were conducted using the FactoMineR and umap R packages, respectively. Statistical analysis was carried out at the peptide level using the ROPECA approach (via the PECA R package), which employs a reproducibility-optimized statistical test (ROTS) tailored for DIA-MS data. Proteomic analyses were conducted at Proteomics Core at Weill Cornell Medicine Qatar (WQM-Q).

### TPL1 amplification and In vitro transcription

To clone TPL1 transcript (ENST00000702307) Total RNA was reverse transcribed using the High-Capacity cDNA Reverse Transcription Kit (Applied Biosystems, Foster City, CA, USA) following the manufacturer’s instructions. The resulting cDNA was used as a template for semi-quantitative PCR using the KAPA Taq DNA Polymerase Kit (Kapa Biosystems, Wilmington, MA, USA) and TPL1 specific primers (forward primer containing a T7 promoter, Table S2). The PCR amplicon was analyzed by electrophoresis on a 1.5% agarose gel stained with ethidium bromide and visualized using a gel documentation system. PCR product (900 bp) was cloned using the pCR™2.1vector TA Cloning Kit (Invitrogen, USA) according to the manufacturer’s protocol. Following ligation, 2 μL of the reaction was used to transform chemically competent E. coli DH5α cells followed by plasmid isolation, PCR validation and Sanger sequencing verification (Supplementary datafile 1) using Applied Biosystems 3130xl Genetic Analyzer.

In vitro transcription was performed using the HiScribe™ T7 Quick High Yield RNA Synthesis Kit (NEB, #E2050S, Ipswich, MA, USA) following the manufacturer’s protocol, with modifications to incorporate fluorescently labeled nucleotides. The DNA template (PCR product) containing a T7 promoter upstream of the insert was purified using PureLink™ Quick PCR Purification Kit (Invitrogen, USA). To generate Cy3-labeled RNA, Cy3-UTP (APExBIO, USA) was incorporated into the transcription reaction. Each 20 μL reaction contained 5 μL of NTP buffer mix, 5 μL of Cy3-UTP, 1 μg of PCR product, 1 μL of 100 mM DTT, and 2 μL of T7 RNA Polymerase Mix. To remove the DNA template following transcription, 30 μL of nuclease-free water and 2 μL of DNase I were added directly to the reaction, followed by incubation at 37°C for 15 minutes. The Cy3-labeled RNA was subsequently purified using the Monarch^®^ RNA Cleanup Kit (NEB #T2040, MA, USA) according to the manufacturer’s instructions. RNA integrity was evaluated using an Agilent Bioanalyzer, and concentration was quantified using a Qubit fluorometer (Thermo Fisher Scientific, USA).

### TPL1 protein interaction profiling using human proteome microarray

To assess protein interactors of TPL1, we performed protein array hybridization as previously described ^34^. In brief, 50 pmol of Cy3-labeled TPL1 RNA was hybridized to the Human Proteome Microarray (HuProt v4.0, CDI Laboratories), which includes over 21,000 full-length human proteins and isoforms. Cy3-tagged firefly luciferase RNA was used as a negative control to eliminate non-specific interactions and background signal. These experiments were conducted at the HBKU Proteomics Facility (Hamad Bin Khalifa University, Qatar). Hits were identified using a defined analysis pipeline (CDI labs). A protein spot was considered a positive interactor if it exhibited a signal at least four-fold higher (log2 > 2) than the control-only median for that spot and had a Z-score greater than 3 within the individual sample array. This dual threshold, log2 fold change and Z-score, helps eliminate artifacts caused by background staining variability and ensures that detected signals represent true outliers within each array. Downstream analyses were based on binary hit status (presence or absence of interaction), semi-continuous Z-score distributions for all passing hits, or directly on raw log₂-transformed median fluorescence intensity (MFI) data. The gene effect scores for the identified TPL1 binding proteins were retrieved from the genome-wide CRISPR-Cas9 functional screen data from the Achilles project in TNBC models as described before ^35^.

### Identification of potential TPL1-binding miRNAs and their gene targets

Small RNA libraries were prepared from 100 ng of total RNA using the QIAseq miRNA NGS workflow (QIAGEN, Hilden, Germany). The protocol involved sequential 3′ and 5′ adaptor ligations, followed by reverse transcription and cDNA library amplification using unique tube indices (QIAseq miRNA NGS 96 Index IL kit). Library quality and concentration were assessed using the Qubit dsDNA HS assay (Invitrogen), and fragment size distribution was evaluated with the Agilent 2100 Bioanalyzer using a DNA1000 chip (Agilent Technologies). Libraries were pooled and sequenced on an Illumina platform.

Sequencing data were processed by mapping FASTQ files to the miRBase v22 database using the built-in small RNA analysis workflow in CLC Genomics Workbench v24.0.2. miRNA expression levels were estimated as total counts, normalized to counts per million (CPM), CLC Genomics Workbench v24.0.2. Differential expression analysis was then performed to identify miRNAs significantly upregulated in TPL1 knockdown cells using AltAnalyze v.2.1.3 as we described before ^36,37^.

To explore potential molecular interactions, the upregulated miRNAs were screened for predicted binding to TPL1 using the miRanda algorithm (version 3.3a) ^38^. In parallel, mRNAs that were downregulated upon TPL1 knockdown were identified from RNA-seq data. These datasets were integrated using the MicroRNA Target Filter module in Ingenuity Pathway Analysis (IPA, Qiagen), allowing identification of high-confidence or experimentally validated mRNA targets of the TPL1-associated miRNAs. The resulting data were used to construct a TPL1–miRNA–mRNA regulatory network, as previously described ^39^.

## Results

### Robust functional profiling of TNBC-associated lncRNAs employing CRISPR-del screen

To generate a comprehensive lncRNA dependency map in TNBC, we employed a high-throughput dual-excision CRISPR-del system (Figure 1A). This strategy leverages loss-of-function (LOF) perturbations by targeting the TSS of lncRNA genes using pgRNAs, effectively suppressing lncRNA expression. This approach offers several advantages over full-gene deletion, ensuring uniform deletions, enhancing efficiency, and reducing the risk of false positives caused by inadvertent deletion of overlapping genomic regulatory elements.

**Figure 1.**
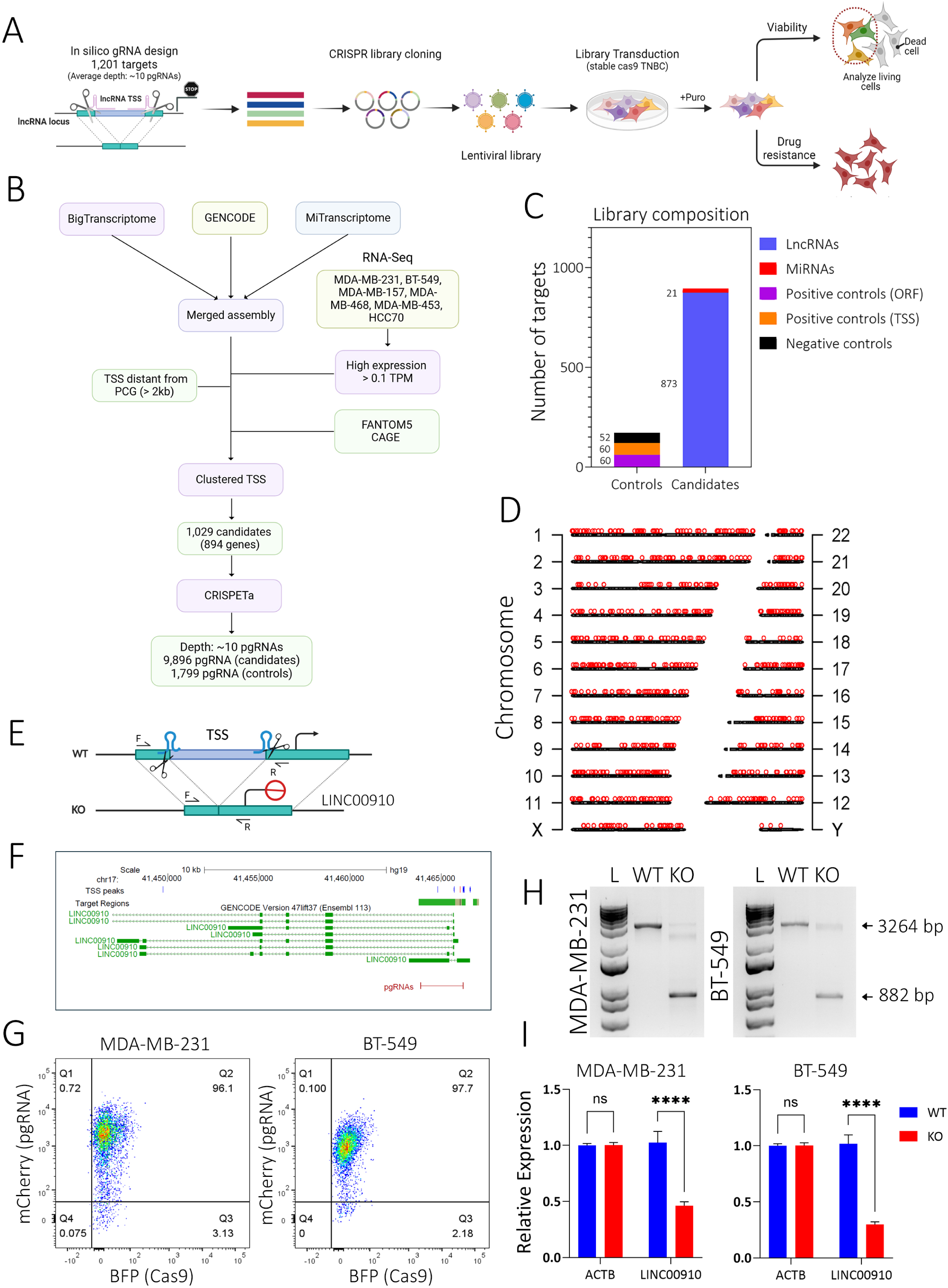
CRISPR-Cas9 deletion screen system for identifying essential and chemosensitive lncRNAs in TNBC. **(A)** Schematic overview of the paired-guide RNA (pgRNA) library design and the CRISPR-deletion functional screening strategy. **(B)** Library design based on a merged annotation of GENCODE v19, MiTranscriptome, and BigTranscriptome, encompassing 1,029 transcription start sites (TSSs). For lncRNAs with bidirectional promoters or overlapping TSSs (≤ 50 bp apart), a single representative TSS was selected. TSS annotations were validated using FANTOM breast and pan-cell line data. **(C)** The library comprised 11,695 pgRNAs, with an average coverage of ∼10 unique pgRNAs per target. Control pgRNAs targeted neutral genomic loci and positive control protein-coding genes (PCGs), including sequences within open reading frames (ORFs) and TSS regions. Overall, the library was composed of 97.6% lncRNA-targeting and 2.4% miRNA-targeting pgRNAs. **(D)** Chromosomal distribution of pgRNAs across all 23 human chromosomes. **(E)** Schematic illustrating the CRISPR-mediated TSS deletion strategy for LINC00910. **(F)** Genomic map showing LINC00910 coordinates and the positions of pgRNAs used for its TSS deletion. **(G)** Dot plot from flow cytometry analysis confirming successful transduction of MDA-MB-231 and BT-549 TNBC cell lines with Cas9 (BFP, x-axis) and pgRNAs (mCherry, y-axis). **(H)** Genomic PCR genotyping confirming deletion of the LINC00910 TSS region in knockout (KO) compared to wild-type (WT) MDA-MB-231 and BT-549 cells. **(I)** RT-qPCR analysis validating LINC00910 knockdown in KO MDA-MB-231 and BT-549 cells. Data are presented as mean ± S.D., n = 6. ns, not significant; ****p < 0.00001.

Firstly, we constructed a customized screening library that comprehensively covers the ncRNA transcriptome specific to TNBC, utilizing transcriptomic data from TNBC cell models. To minimize false positives from off-target effects, we systematically excluded target regions overlapping protein-coding gene (PCG) transcripts or those within 5 kilobases of the nearest PCG TSS. We also ensured the removal of candidate TSSs near PCG exons, both upstream and downstream. By integrating and filtering data from three resources—GENCODE R19, MiTranscriptome, and BigTranscriptome—we assembled a final set of 1,029 expressed lncRNA (GENCODE-annotated), miRNA, and novel unannotated lncRNA candidate TSSs (Figure 1B). This set was subsequently dubbed ’candidatetnbc 1’ and so forth. In cases where lncRNA candidates featured bidirectional promoters or overlapping TSSs (TSSs <= 50 bp), they were merged into a single TSS. Correct TSS was verified using FANTOM breast and FANTOM all primary and cancer cell lines data. Any lncRNA candidate not annotated in the GENCODE R19 was classified as novel. The candidate distribution in the final library contained 97.6 % lncRNAs, among those 39.4% were annotated in the GENCODE R19 reference, in addition to 2.4% miRNAs (Figure 1C). Additionally, we included pgRNAs designed to target neutral control loci, those not expected to exert any influence on cellular phenotype. Furthermore, we incorporated positive control PCGs based on previous genomic screens with well-established roles in cell growth ^35^. These lncRNA target genes served as the foundation for our CRISPR-del library, which features a depth of ∼10 unique pgRNAs per target, comprising 11,695 pgRNAs (Figure 1C and Table S3). Chromosomal distribution of pgRNAs across all 23 human chromosomes are shown in Figure 1D. The individual sgRNAs and pgRNAs scores are provided in Figure S2. Upon cloning into lentiviral vector backbone, the pgRNA library was built from a total of 3.6 x 10^7^ single colonies which covered the library for more than 1,000 folds. A 150 bp paired-end sequencing identified a total of 11,686 gRNA pairs out of the 11,695 designed gRNA pairs (99.92% coverage). The alignment results are shown in Table S4, while the nucleotide distribution at each position is summarized in Table S5.

To broaden the relevance of our findings to TNBC, we conducted parallel screens using two well-established TNBC models, MDA-MB-231 and BT-549. First, we generated cell lines with stable high-level Cas9 expression. To validate suitability of our system for pgRNA CRISPR-del screening, we used selected pgRNAs targeting the LINC00910 TSS from our library (candidate 256). A schematic of the pgRNAs targeting the LINC00910 TSS in the Double Excision CRISPR Knockout vector (pDECKO) is shown in Figure 1E. Genomic map showing LINC00910 coordinates and the positions of pgRNAs used for its TSS deletion are shown in figure 1F. Flow cytometry analysis showed > 96% co-expression of Cas9 (BFP) and LINC00910 pgRNAs (mCherry) in MDA-MB-231 and BT-549 (Figure 1G) TNBC models. Genotyping of the targeted TSS confirmed successful deletion in knockout (KO) versus wild-type (WT) MDA-MB-231 and BT-549 models (Figure 1H). RT-qPCR further verified significant suppression of LINC00910 expression in KO MDA-MB-231 and BT-549 cells, demonstrating the efficacy of CRISPR-del for lncRNA loss-of-function (Figure 1I).

### CRISPR-del screens revealed lncRNA dependencies and sensitivity to doxorubicin in TNBC

We adapted a pooled CRISPR-del screening approach to identify lncRNAs essential for TNBC cell proliferation and chemosensitivity through a negative selection screen, where targets’ pgRNAs become depleted (“dropout”). Figure 2A illustrates the CRISPR-del genetic screening strategy and analysis. Transducing TNBC cells with the lentiviral library achieved ∼98% (MDA-MB-231) and ∼97% (BT-549) enrichment after blasticidin selection at T0 (Figure 2B and 2C, respectively). The extracted genomic DNA was then amplified with primers spanning the pgRNAs and NGS data were subsequently analyzed, revealing over 60% alignment (Figure S3) and high correlation between biological replicates at the gene-level log2 fold change (Figure 2D). Drop out assay data revealed strong correlation (R2= 0.88) between the MDA-MB-231 and BT-549 models (Figure 2E).

**Figure 2.**
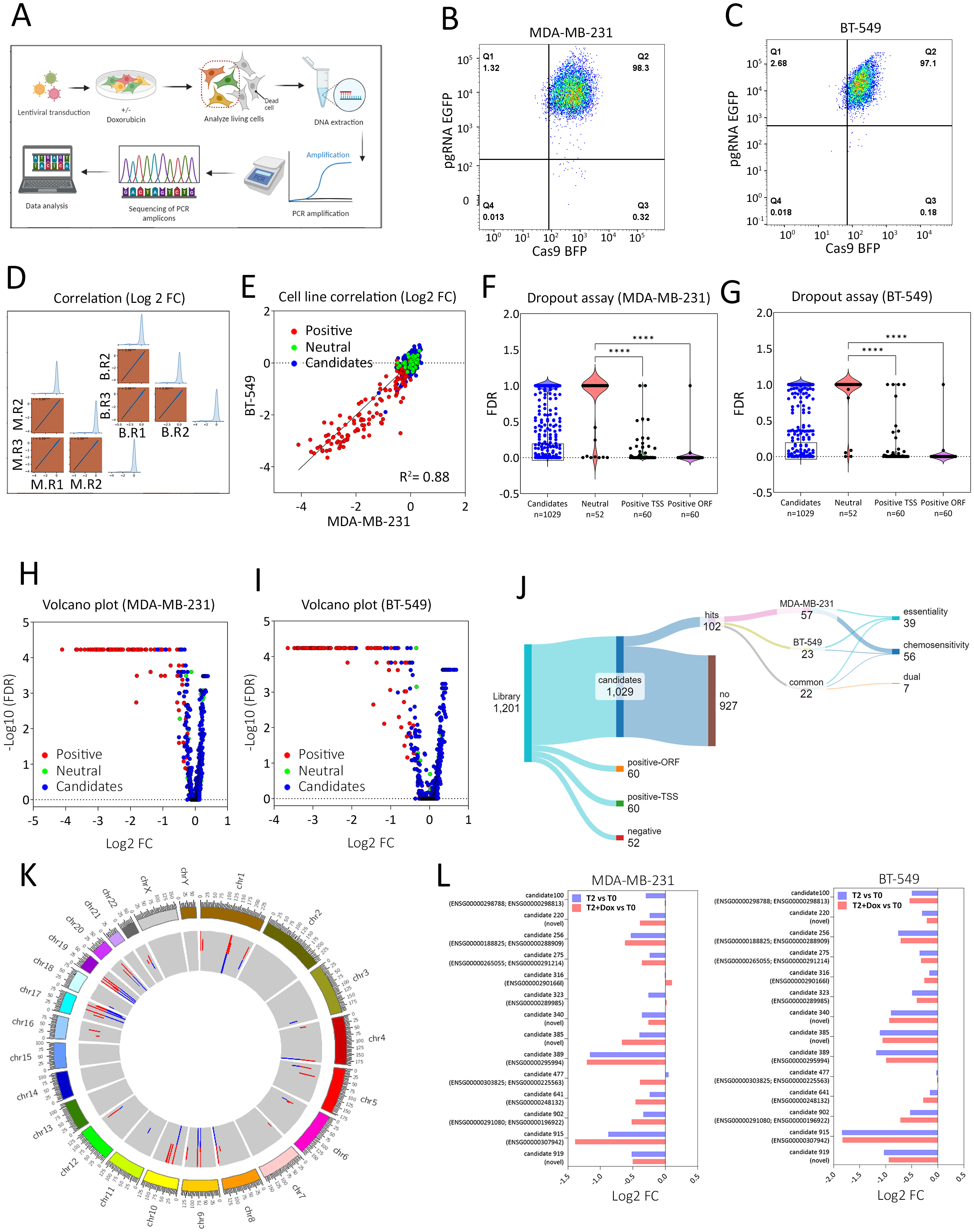
Identification of candidate lncRNAs essential for TNBC survival and chemosensitization. **(A)** Schematic presentation of the CRISPR-del screening strategy. Flow cytometry depicting the percentage of double positive (Cas9 (BFP) and pgRNA (EGFP)) in the MDA-MB-231 **(B)** and BT-549 **(C)** TNBC models after puromycin selection (T0). **(D)** Gene level log2 FC correlation from the dropout assay in MDA-MB-231 (M) and BT-549 (B) models. **(E)** Log2 FC correlation from the drip out assay between MDA-MB-231 (x-axis) and BT-549 (y-axis) depicting the positive and neutral controls, in addition to candidates. P-value distribution of lncRNA candidates and positive and negative controls from the MDA-MB-231 **(F)** and BT-549 **(G)** dropout screen. Volcano plot for MDA-MB-231 **(H)** and BT-549 **(I)** illustrating identified hits from the screen. Positive (red), neutral (light green), and candidates (blue). **(J)** Sankey diagram illustrating library composition and the identified hits in MDA-MB-231 and BT-549 TNBC models, categorized as essentiality, chemosensitivity, or dual. Each screen was done in triplicate. **(K)** Circos plot illustrates the chromosomal distribution of identified hits from the screen. **(L)** Log2 FC for the fourteen commonly identified essential hits in MDA-MB-231 and BT-549 and the added toxicity from combination with doxorubicin based on CRISPR-del screen.

As expected, pgRNAs targeting positive-control genes were significantly depleted in the dropout screens, while neutral controls were not in the MDA-MB-231 (Figure 2F) and BT-549 (Figure 2G) models. To assess the efficacy of promoter deletion, we compared pgRNAs targeting PCGs with conventional ORF mutations and promoter deletions, the latter mirroring our TSS-targeting strategy for lncRNAs. Promoter-deletion pgRNAs showed a discernible but attenuated phenotypic impact, indicating that CRISPR-del screens targeting TSSs are inherently less effective than those targeting ORFs (Figure 2F and 2G) consistent with previous report ^14^. Top affected candidates from the MDA-MB-231 (Figure 2H) and BT-549 (Figure 2I) screen are depicted as volcano plots. Library composition and the identified hits in MDA-MB-231 and BT-549 TNBC models are summarized as sankey diagram, revealing a total of 102 identified hits in both models under proliferation and doxorubicin conditions (Figure 2J and Table S6). The chromosomal distribution of all identified hits from the screen are illustrated as circos plot (Figure 2K). Our proliferation screen identified 14 common hits from both models shown as log2 fold changes either as single perturbations or in combination with doxorubicin (Figure 2L). Intersecting these 14 candidate TSSs with the GENCODE version 47lift37 (Ensembl 111) GTF revealed 4 novel candidates, while existing annotations were found for the remaining 10 candidates (Figure 2L and Table S7).

### Functional validation of candidate lncRNAs TPL1 and PROLIT1 using ASO-mediated suppression

Based on our CRISPR-deletion screen, candidate 323 (named TNBC Promoting LncRNA 1 [TPL1]) and candidate 389 (named Proliferation LncRNA in TNBC 1 [PROLIT1]) emerged as potential functional lncRNAs in TNBC. The genomic coordinates of both lncRNAs, along with the set of paired guide RNAs (pgRNAs) used to target their TSSs in the screen are shown in Figure 3A. TPL1 corresponds to Ensembl gene ENSG00000289985, while PROLIT1 corresponds to ENSG00000305847. The results from the dropout screen revealed significant reduction in pgRNA abundance targeting the TSSs of TPL1 and PROLIT1 in MDA-MB-231 and BT-549 cells at T2, suggesting a selective disadvantage upon their loss (Figure 3B) in the proliferation screen. To functionally validate these findings, we employed antisense oligonucleotide (ASO)-mediated knockdown approaches targeting TPL1 and PROLIT1 transcripts (Figure 3C). Two independent ASOs were used per lncRNA, and their effects were assessed in MDA-MB-231 cells. Colony formation assays showed a robust reduction in clonogenic potential upon ASO transfection compared to non-targeting control (NC) ASO, with remarkable decreases in cell viability (Figure 3D, upper panel) and effective suppression of TPL1 and PROLIT1 expression (Figure 3D, lower panel). Dose-response analysis across a range of ASO concentrations (0, 7.5, 15, and 30 nM) further confirmed the inhibitory effects of TPL1 and PROLIT1 suppression in both MDA-MB-231 and BT-549 cell lines (Figures 3E and 3F, respectively). The calculated IC50 values indicated dose-dependent sensitivity (dotted lines). Concordantly, quantitative proliferation analysis using the Operetta CLS High-Content Analysis System demonstrated a marked decrease in cell proliferation following ASO-mediated knockdown in MDA-MB-231 cells, which maximized at 96 hr (Figure 3G).

**Figure 3.**
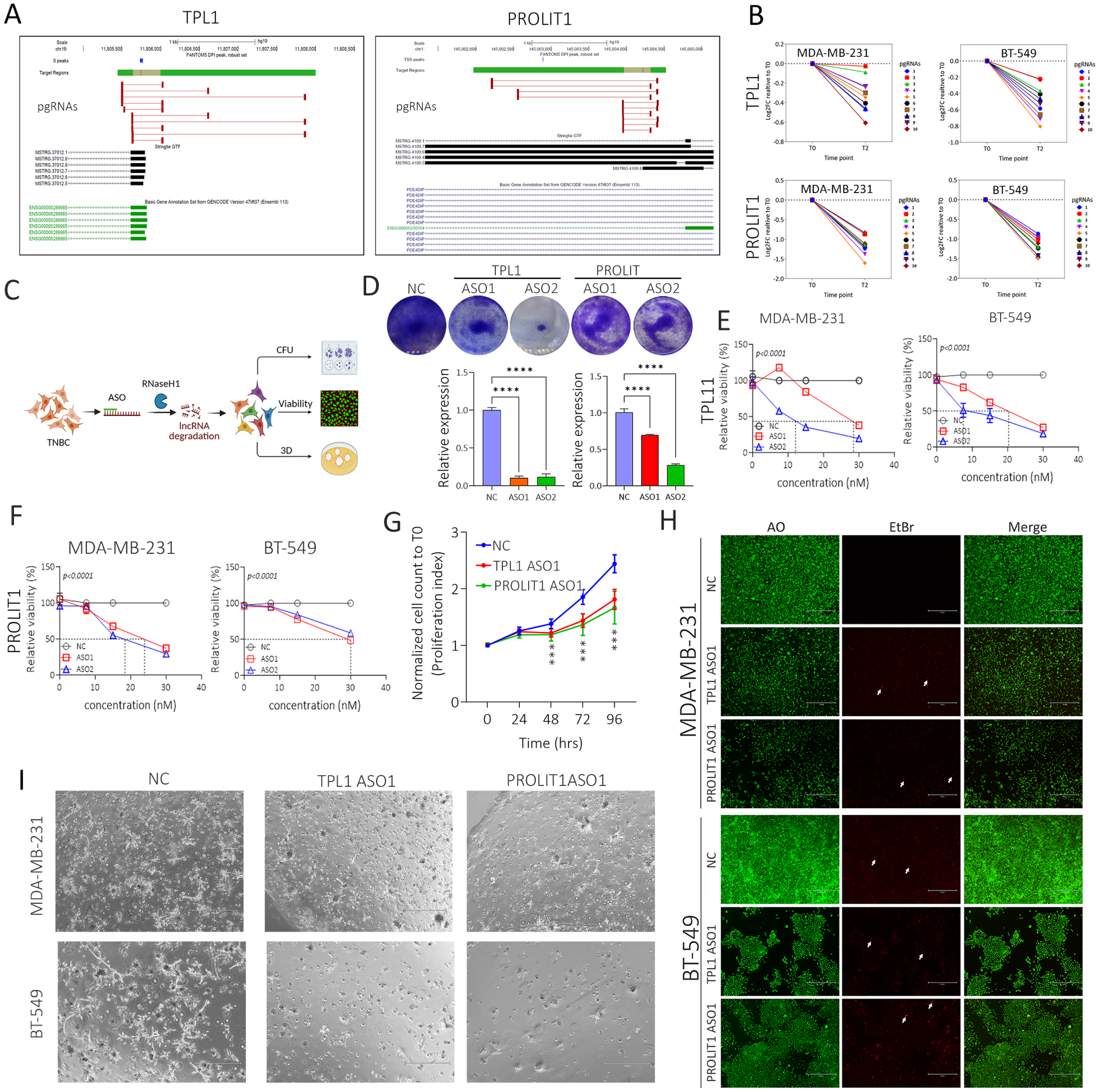
Functional validation of candidate lncRNAs TPL1 and PROLIT1 using CRISPR-del and ASO-based approaches. **(A)** Genomic loci of candidate 323 (TPL1) and candidate 389 (PROLIT1), along with the set of pgRNAs targeting their transcription start sites (TSSs) used in the CRISPR-deletion screen. Overlapping gene transcripts based on GENCODE v47 are shown. **(B)** Guide plots showing the log₂ fold change (log2 FC) in pgRNA abundance targeting TPL1 and PROLIT1 TSSs at T0 and T2 (after 21 population doublings) in MDA-MB-231 and BT-549 TNBC cell lines. **(C)** Schematic representation of the experimental strategy for functional validation using antisense oligonucleotides (ASOs). **(D)** Representative images from colony formation (CFU) assays demonstrating the effects of two independent ASOs targeting TPL1 and PROLIT1 in MDA-MB-231 cells. Efficacy of TPL1 and PROLIT1 knockdown using each ASO is shown below. Data are presented as mean ± S.E.M. Dose–response curves for ASO-mediated suppression of TPL1 **(E)** and PROLIT1 **(F)** in MDA-MB-231 and BT-549 cells, treated with 0, 7.5, 15, and 30 nM ASOs. Data are presented as mean ± S.D., n = 3. Statistical significance is indicated on each plot; dotted lines represent estimated IC50 values. **(G)** Proliferation curves generated using the Operetta CLS High-Content Analysis System, showing quantitative proliferation of MDA-MB-231 cells treated with ASOs targeting TPL1 and PROLIT1. **(H)** Representative live (green) and dead (red) staining of MDA-MB-231 and BT-549 TNBC cells on day 5 post suppression of the indicated candidates using ASOs. NC ASO was used for comparison. Images were taken using an Olympus IX73 fluorescence microscope (Olympus, Tokyo, Japan) (10X, Scale bar = 100uM). Two arrows highlight repetitive cells staining with EtBr (red) in KD cells. **(I)** Phase-contrast images showing inhibition of 3D organoid formation in MDA-MB-231 (upper panel) and BT-549 (lower panel) TNBC models following suppression of TPL1 and PROLIT1 using the indicated ASOs compared to NC ASO. Images were captured on day 7 using the EVOS Cell Imaging System (10× magnification, scale bar = 1000 µm).

Live/dead cell staining on day 5 post-ASO-mediated suppression of the indicated candidates revealed reduction in cell proliferation and increased cell death in MDA-MB-231 and BT-549 TNBC cells compared to the negative control ASO (Figure 3H). Furthermore, the suppression of TPL1 and PROLIT1 disrupted 3D organoid formation in both MDA-MB-231 (Figure 3I, upper panel) and BT-549 (Figure 3I, lower panel) models, as shown by phase-contrast microscopy on day 7 post-transfection. These results support a functional role for TPL1 and PROLIT1 in promoting TNBC cell proliferation and 3D growth.

### ASO-mediated suppression of TPL1 and PROLIT1 impairs TNBC cell migration and invasion

To investigate the functional role of TPL1 and PROLIT1 in TNBC cell motility hallmark, we performed migration and invasion assays following ASO-mediated knockdown in MDA-MB-231 cells. The experimental workflow is illustrated in Figure 4A. Wound healing assays were conducted using EGFP-labeled MDA-MB-231 cells, with gap closure monitored at 0, 24, and 36 hr post scratching. Our data revealed delayed wound closure upon suppression of PROLIT1, but not TPL1 compared to the control ASO (Figure 4B). Quantitative analysis showed a significant reduction in gap closure over time in PROLIT1 ASO-treated cells (Figure 4C), with a marked increase in residual wound area at 36 hr (Figure 4D).

**Figure 4.**
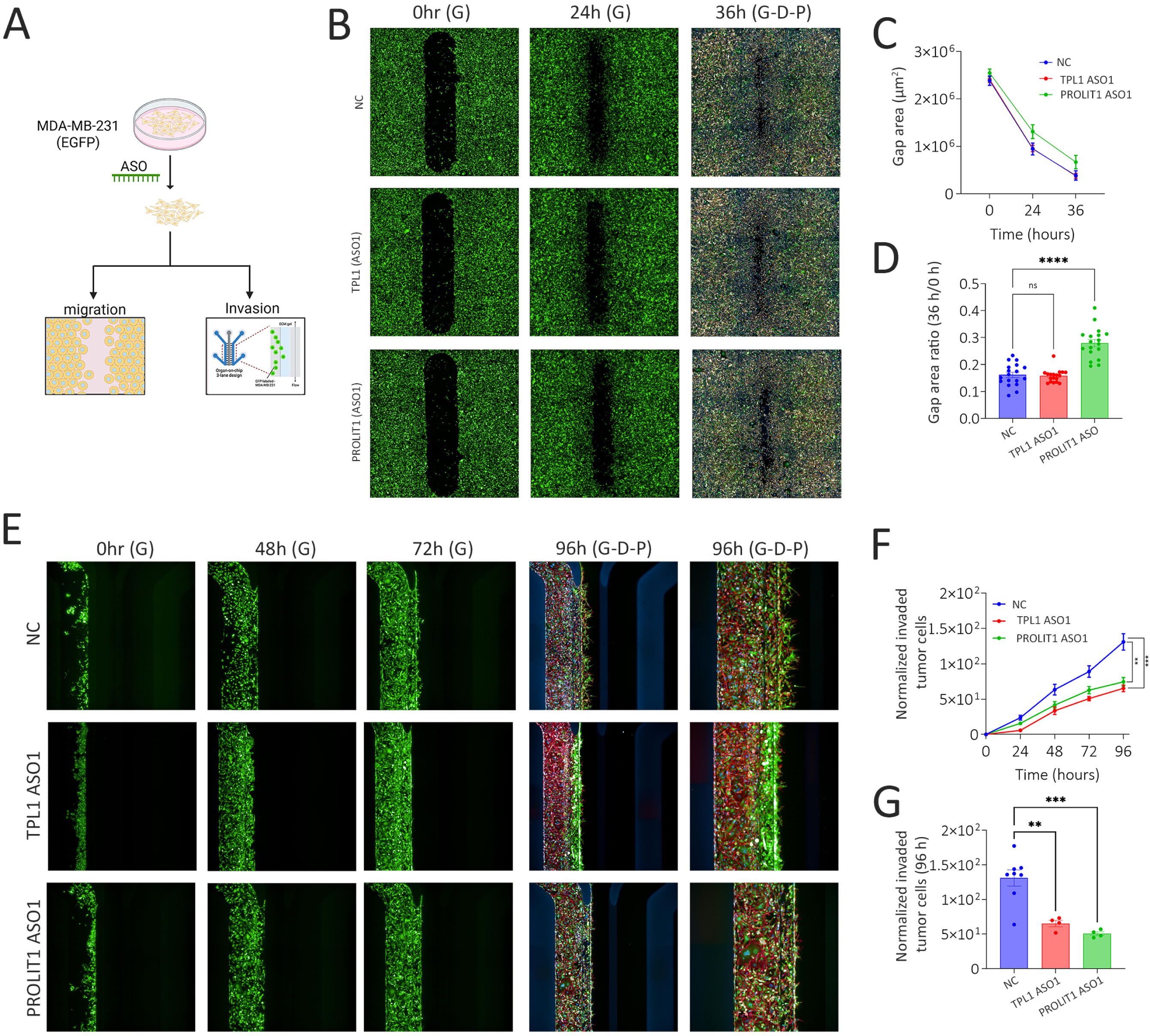
Suppression of cell migration and invasion by ASO-mediated targeting of TPL1 and PROLIT1. **(A)** Schematic illustrating the strategy used to assess the role of TPL1 and PROLIT1 in regulating TNBC cell migration and invasion using the Operetta CLS High-Content Analysis System. **(B)** Representative images of wound healing (gap closure) in MDA-MB-231 (EGFP-labeled) cells at 0, 24, and 36 hours after scratching. Cells were stained with DAPI (blue), phalloidin (red), and EGFP (green). **(C)** Quantification of the wound gap area under the indicated conditions at 0, 24, and 36 hours after scratching. **(D)** Quantification of gap area at 36 h after scratching, under the indicated conditions. Data are presented as mean ± SEM, n = 18. **(E)** Organ-on-chip invasion assay showing the number of MDA-MB-231 cells invading the extracellular matrix (ECM) at 0, 48, 72, and 96 hours after seeding. The final panel shows a magnified image of each condition at 96 hours. Cells are visualized using EGFP (green), DAPI (blue), and phalloidin (red). **(F)** Quantification of the normalized number of invaded cells at 0, 48, 72, and 96 hours after seeding. **(G)** Quantification of normalized invaded MDA-MB-231 cells at 96 hours post-ASO transfection. Data are presented as mean ± SEM, n = 4. p < 0.005, *p < 0.0005.

To further evaluate the role of both lncRNAs in tumor cell invasion, we utilized a 3D organ-on-chip invasion assay. MDA-MB-231 cells transfected with either TPL1 or PROLIT1 targeting ASOs exhibited reduced invasion into the extracellular matrix (ECM) over a 96-hour period. Images acquired at 0, 48, 72, and 96 hours after seeding confirmed a substantial decrease in ECM invasion upon lncRNA suppression, with fewer cells penetrating the matrix compared to control cells (Figure 4E). Quantification of invading cells showed significant reductions in invasion over time, with the most pronounced effects observed at 96 hours (Figures 4F and 4G). These findings demonstrate that PROLIT1 contributes to the migratory, while TPL1 and PROLIT1 modulates invasive properties of TNBC cells, further supporting their functional relevance in promoting tumor progression.

### TPL1 is upregulated in TNBC and regulates multiple cancer hallmarks

The remaining experiments were focused on TPL1, given its novelty and relevance to TNBC. We first examined the expression of TPL1 in TNBC tumors compared to adjacent normal breast tissues using transcriptomic data. TPL1 was significantly upregulated in TNBC (n = 360) relative to adjacent normal tissues (n = 88) (Figure 5A). Further stratification of TNBC cases revealed subtype-specific differences, with the highest TPL1 expression observed in the aggressive basal-like immunosuppressed (BLIS) subtype, followed by immunomodulatory (IM), mesenchymal (MES), and luminal androgen receptor (LAR) subtypes (Figure 5B). These differences were statistically significant (one-way ANOVA, p < 0.0001), suggesting subtype-dependent expression patterns.

**Figure 5.**
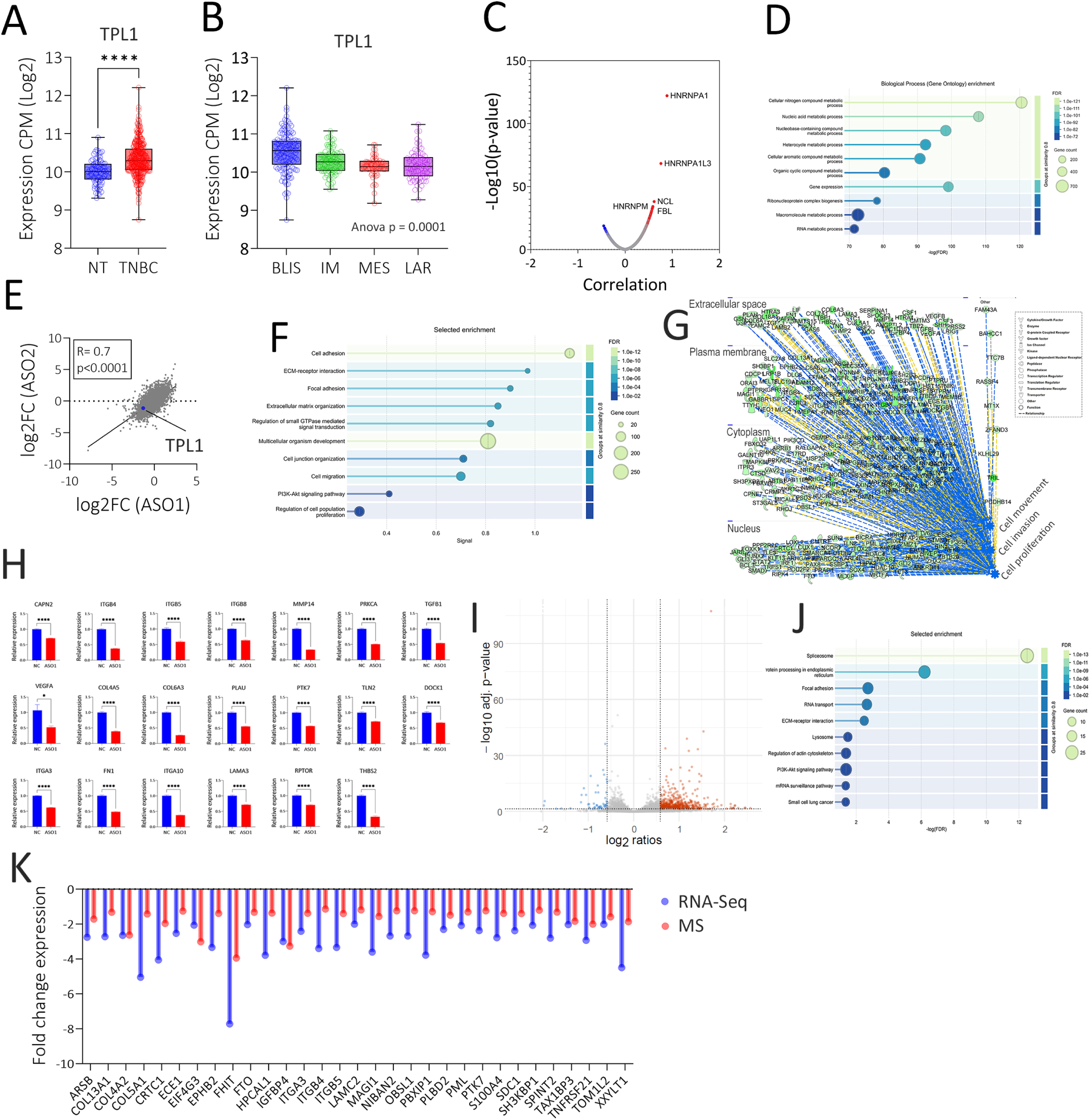
TPL1 is upregulated in TNBC and regulates multiple cancer hallmarks. **(A)** Expression of TPL1 in TNBC tumors (n = 360) versus adjacent normal breast tissues (n = 88). ****p < 0.00005. **(B)** TPL1 expression across TNBC subtypes: basal-like immunosuppressed (BLIS), immunomodulatory (IM), mesenchymal (MES), and luminal androgen receptor (LAR). One-way ANOVA, p < 0.0001. **(C)** Volcano plot showing protein-coding genes correlated with TPL1 expression in TNBC (n = 360). The x-axis represents Pearson correlation values, and the y-axis shows –log10(p) values. The top five positively correlated genes are highlighted. (**D)** Gene Ontology (GO) enrichment analysis of protein-coding genes positively correlated with TPL1 (Pearson r > 0.25, p < 0.05), indicating enrichment in pathways relevant to cancer biology. **(E)** Scatter plot showing the log₂ fold change in gene expression in MDA-MB-231 cells transfected with ASO1 versus ASO2 targeting TPL1. Pearson correlation coefficient (r) and p-value are indicated. **(F)** Enriched functional categories among downregulated genes in TPL1-depleted MDA-MB-231 cells based on STRING protein–protein interaction (PPI) network analysis. **(G)** Network analysis (Ingenuity Pathway Analysis, IPA) of downregulated genes in TPL1-depleted MDA-MB-231 cells, highlighting roles in cancer hallmarks such as proliferation, migration, and invasion. **(H)** RT-qPCR validation of selected downregulated genes identified by RNA-seq, particularly those involved in cell migration, FAK, and AKT signaling pathways, in TPL1-depleted MDA-MB-231 cells. Data are presented as mean ± SEM, n = 6. p < 0.05, ****p < 0.00005. **(I)** Volcano plot showing differentially expressed proteins in TPL1-depleted versus NC ASO-transfected TNBC cells. Top up- and downregulated proteins are labeled on the plot. **(J)** STRING enrichment analysis of downregulated proteins in TPL1-depleted TNBC cells, revealing functional categories and interaction networks. **(K)** Fold-change comparison in expression levels of 31 commonly downregulated genes identified by both RNA-seq (blue bars) and DIA-MS (red bars). Protein identifiers are shown on the x-axis, and log₂ fold change values are shown on the y-axis.

To gain insight into the potential biological functions of TPL1, we performed correlation analysis between TPL1 and all PCGs in TNBC tumors (n = 360). The volcano plot presented in figure 5C highlighted the genes with positive (red) or negative (blue) correlations with TPL1 expression, with the top five positively correlated genes annotated (Figure 5C). The full results from correlation analysis are provided in Table S8. Notably, three of the top genes positively correlated with TPL1 expression, HNRNPA1, HNRNPA1L3, and HNRNPM, are members of the heterogeneous nuclear ribonucleoprotein (hnRNP) family, while several others, including SRSF1, SFPQ, NONO, and FUS, are known RNA-binding and splicing factors. This pattern suggests a possible nuclear function for TPL1, potentially through interactions with the RNA processing machinery. Supporting this hypothesis, analysis of CLIP-Seq data revealed enrichment of TPL1 transcripts on various hnRNP family proteins, in contrast to RBFOX2 RNA-binding protein (Figure S4), concordant with our correlation analysis (Table S8). Gene Ontology (GO) enrichment analysis of genes positively correlated with TPL1 (Pearson r > 0.25, p < 0.05) revealed significant enrichment in pathways associated with nucleic acid metabolic processes, Ribonucleoprotein complex biogenesis, and RNA metabolic processes (Figure 5D and Table S9).

To investigate the transcriptional alterations in TNBC cells upon TPL1 suppression, MDA-MB-231 cells were transfected with two independent ASOs targeting TPL1. RNA-seq analysis comparing gene expression profiles in ASO1 versus ASO2 treated cells revealed strong concordance between the two datasets (Figure 5E; Pearson R= 0.7), supporting reproducibility of the knockdown effects by the two ASOs. STRING-based protein–protein interaction (PPI) analysis of downregulated genes upon TPL1 depletion indicated enrichment in biological processes associated with key cancer-related functions, including Cell adhesion, ECM-receptor interaction, Foal adhesion, Regulation of small GTPase mediated signal transduction, Cell migration, and Cell proliferation (Figure 5F and Table S10). Ingenuity Pathway Analysis (IPA) further revealed that TPL1-regulated genes were involved in critical cancer hallmarks such as tumor cell proliferation, movement, and invasion (Figure 5G and Table S11). Validation by RT-qPCR confirmed the downregulation of several key target genes involved in focal adhesion kinase (FAK), AKT signaling pathways, and cell motility, upon TPL1 knockdown (Figure 5H).

### Mass spectrometry proteomics profiling identifies key protein networks regulated by TPL1 in TNBC cells

To assess the proteomic impact of TPL1 depletion, DIA-MS was performed on MDA-MB-231 cells treated with TPL1- or control-ASOs (n = 5/group). Significant changes in protein abundance were observed (Figure S5A–B, Table S12), with PCA and correlation analyses confirming clear group separation and strong intra-group consistency (Figures S6–S8), supporting both technical reproducibility and biological specificity. Using 0.05 FDR adjusted p value cut off, and 1.5 FC, we observed 48 downregulated and 502 upregulated proteins (Figure 5I). Focusing on all significantly altered proteins (<0.05 FDR adjusted p value), we identified 469 downregulated and 844 upregulated proteins upon TPL1 depletion (Table S13). STRING enrichment analysis of the downregulated proteome uncovered multiple functionally coherent categories and protein– protein interaction networks, including pathways involved in cytoskeletal regulation, cell migration, and RNA processing (Figure 5J). Integrative analysis comparing RNA-seq and DIA-MS data identified 31 genes commonly downregulated at both transcript and protein levels (Figure 5K). Interestingly, enrichment analysis on those 31 genes further revealed a role in Cell migration, Cell adhesion, ECM-receptor interaction, PI3K-Akt signaling pathway, and Focal adhesion (Table S14). These findings suggest that TPL1 promotes TNBC aggressiveness through regulation of oncogenic signaling pathways and gene networks central to cancer progression, which is concordant with our functional data.

### Proteomic array screening reveals high-confidence protein interactors of TPL1 and suggests roles in RNA regulation and cancer signaling

LncRNAs are known to exert their functions through direct interactions with various RNA-binding proteins ^40^. To identify direct protein interactors of TPL1, we employed a proteomic screening approach using the HUPROT human version 4.0 protein array (Figure 6A). Firefly luciferase RNA was included as a negative control to account for non-specific binding. Analysis of Z-score distribution revealed a clear enrichment of specific interactors, with the top 20 proteins (Z score ≥ 3.0) visualized as a heatmap based on signal intensity relative to the control RNA sample (Figure 6B and 6C). Several proteins, including ACTB, B2M, and TUBA1A, which are not expected to bind RNAs were not enriched in our analysis (Figure 6C). Among the highest-ranking hits were EIF4B, a translation initiation factor; TLE5, a transcriptional corepressor; and TARBP2, a double-stranded RNA-binding protein involved in miRNA processing. Additional notable interactors included NACC1, a known oncogene in breast and ovarian cancer; MDM2, a key regulator of p53; and MAP2K6, a component of the MAPK signaling cascade. STRING enrichment analysis on TPL1-interacting proteins revealed possible interaction between PLEK2, DEPC1, and MTMR2. We next examined Tier 2 interactors (2.5 ≤ Z-score < 3.0), identifying 96 additional proteins that could potentially associate with TPL1 (Figure 6B and Table S15). Several of these were classified as RNA-binding proteins, including HNRNPH1, NSUN4, PABPC1L2A, PABPC3, IGF2BP2, SRP19, MTRES1, NPM1, and RPL12. Interestingly, a subset of these interactors (tier 1 and tier 2)—EIF4B, PFKM, STIP1, RAN, NOLC1, PRDX2, METTL2B, RPL14, NPM1, RPL12, and CACYBP—also exhibited positive transcript-level correlation with TPL1 expression in TNBC patient samples (Table S8). Of particular note, RAN—a GTP-binding nuclear protein and GTPase—was identified as a TPL1 interactor, aligning with our pathway analysis that revealed suppression of GTPase signaling following TPL1 knockdown. To assess the potential biological significance of these interactions, we examined genome-wide CRISPR screening data from DepMap ^35^, identifying tier 1 (EIF4B, CACYBP, MDM2, and TARBP2) and tier 2 (RAN, RPL12, RPL14, SNAPC2, PSMD4, WARS1, SRP19, UBA2, NPM1, HNRNPH1, HMGN2, NOLC1, NSUN4, HSD17B10, TWF1, TMX2, and PRDX2) as top dependency genes, highlighting their functional relevance in TPL1-mediated regulation (Figure 6D). These findings suggest that TPL1 functionally associate with regulators of translation, transcriptional repression, RNA metabolism, and stress signaling pathways.

**Figure 6.**
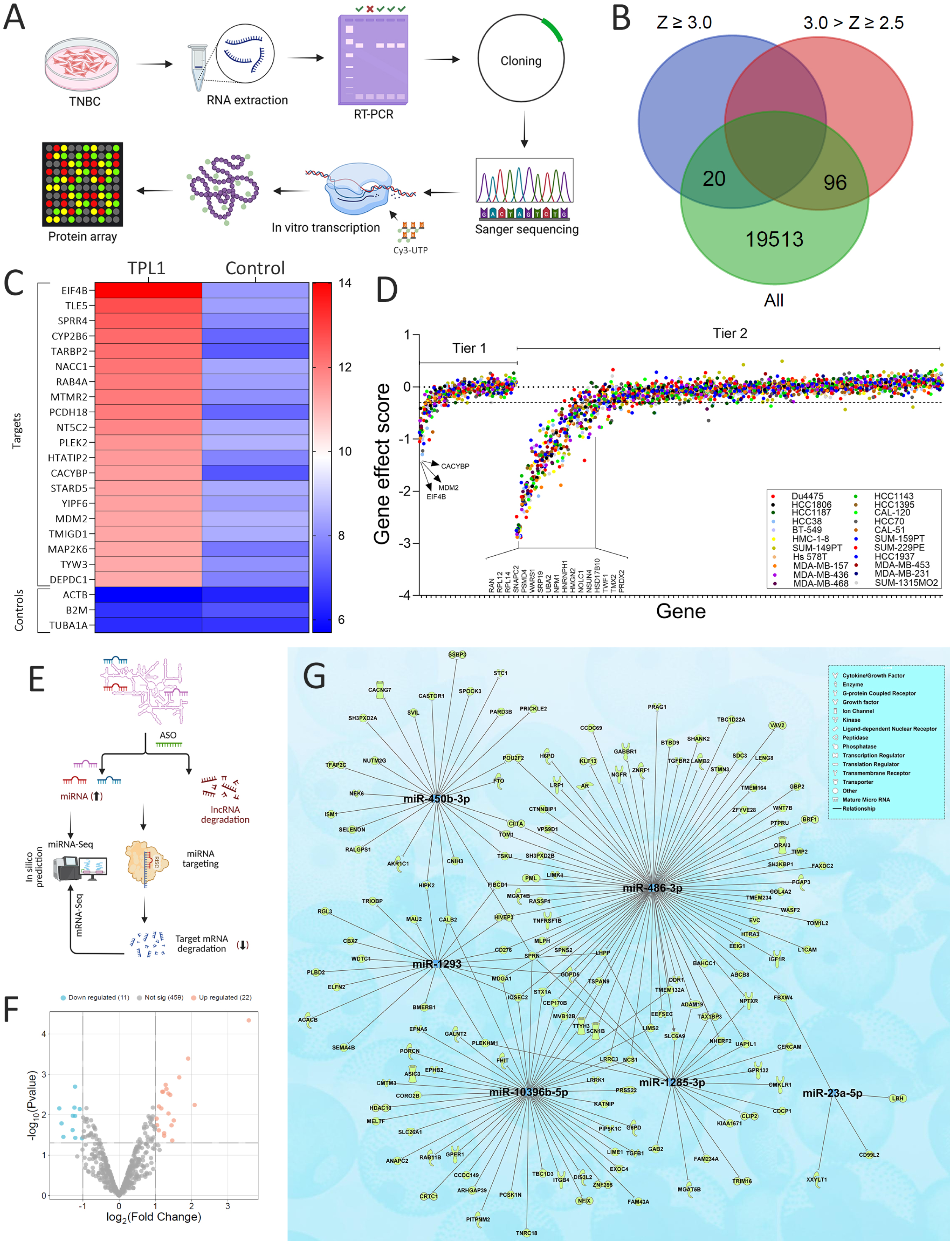
Identification of proteins interacting with TPL1 using proteomic arrays. **(A)** Schematic representation of the experimental strategy used to identify proteins that directly interact with TPL1. **(B)** Venn diagram illustrating the distribution of identified protein hits based on Z-score thresholds: Z ≥ 3.0 and 2.0 < Z ≤ 3.0. **(C)** Heatmap showing 20 tier 1 protein hits (Z ≥ 3.0) identified from the array. The color scale represents signal intensity, comparing TPL1-hybridized arrays to the control RNA sample**. (D)** Scatter plot showing the dependency of TPL1-interacting proteins across TNBC cell lines. Gene effect scores from DepMap CRISPR screens are displayed for the (tier 1 Z ≥ 3.0) and tier 2 (2.0 < Z ≤ 3.0) TPL1-binding proteins in 22 TNBC models. Each dot represents the gene effect score in a single cell line. **(E)** Schema depicting the approach used to identify miRNAs sponged by TPL1 and their regulatory mRNA network. **(F)** Volcano plot showing differentially expressed miRNAs in TPL1 knockdown MDA-MB-231 cells. The plot displays log₂ fold change versus –log₁₀(p-value) for miRNAs detected by small RNA-seq following TPL1 knockdown. Light green dots represent significantly upregulated miRNAs, and light blue dots represent significantly downregulated miRNAs, based on a threshold of ≥2.0-fold change and p < 0.05. **(G)** Network plot illustrating the proposed miRNA–mRNA regulatory interactions potentially mediated by TPL1 in TNBC. Five miRNAs were identified that are (i) significantly upregulated in TPL1-knockdown cells, and (ii) predicted to bind TPL1 based on miRanda v3.3a analysis (Score ≥140, Energy ≤ –50 kcal/mol). To further define downstream regulatory effects, Ingenuity Pathway Analysis (IPA) microRNA Target Filter was used to identify high-confidence or experimentally validated target genes of these miRNAs. These targets were subsequently cross-referenced with RNA-seq data to retain only genes that were downregulated in TPL1-depleted TNBC cells, supporting a ceRNA-like mechanism in which TPL1 may act as a miRNA sponge. The final network includes miRNAs and their filtered targets, highlighting potential oncogenic or tumor-suppressive nodes affected by TPL1 loss.

LncRNAs are increasingly recognized as key regulators of gene expression, often functioning as molecular sponges that sequester specific miRNAs, thereby relieving their target mRNAs from miRNA-mediated repression ^41^. Schema depicting the approach used to identify miRNAs sponged by TPL1 and their regulatory mRNA network is shown in figure 6E. To explore the potential of TPL1 to act as a miRNA sponge in TNBC, we performed small RNA sequencing following TPL1 knockdown in MDA-MB-231 cells. This analysis identified 22 significantly upregulated miRNAs (≥2.0-fold change, p < 0.05; Figure 6F and Table S16). We next used miRanda (v3.3a) to predict miRNA-TPL1 interactions and identified seven miRNAs, hsa-miR-486-3p, hsa-miR-10396b-5p, hsa-miR-450a-2-3p, hsa-miR-1293, hsa-miR-1285-3p, hsa-miR-10396b-5p, hsa-miR-1246, and hsa-miR-23b-5p, as having strong binding potential to TPL1 transcripts (score ≥140 and binding energy ≤ –25 kcal/mol; Table S17). In contrast, none of the 11 downregulated miRNAs in TPL1 KD cells met this threshold, indicating minimal or no predicted interaction with TPL1 (Table S18). Fisher’s exact test revealed a potential enrichment of TPL1-binding sites among upregulated miRNAs (p = 0.07; odds ratio = inf), suggesting that TPL1 preferentially interacts with and represses a subset of miRNAs that become de-repressed upon its silencing.

Next, we applied the IPA microRNA Target Filter to retrieve experimentally validated and/or predicted targets (TargetScan Human) of these miRNAs and cross-referenced them with genes downregulated in TPL1-KD cells from our transcriptome data. Notably, miR-10396b-5p, miR-1285-3p, miR-1293, miR-23b-5p, miR-450a-2-3p, miR-486-3p, and miR-7977 were found to regulate multiple transcripts involved in key cellular processes, including signal transduction (e.g., IQSEC2, HIPK2), cell adhesion and migration (e.g., ITGB4, CDCP1, LIMS2), and transcriptional regulation (e.g., TGFB1, TGFBR2, CIITA) (Figure 6G and Table S19). Several genes, including SPRN, MDGA1, LHPP, and TALB2, were common targets of more than one miRNA, suggesting coordinated post-transcriptional regulation.

Collectively, these findings support a model wherein TPL1 functions as ceRNA that sponges tumor-suppressive miRNAs, thereby sustaining the expression of their gene targets. Figure 7 provides Schematic illustration of the dual oncogenic functions of TPL1 in TNBC which function as a molecular scaffold by (Figure 7A) binding to key RNA-binding proteins (RBPs) such as EIF4B, TLE5, and MDM2, thereby potentially modulating transcriptional and post-transcriptional gene expression programs. Concurrently, TPL1 functions as ceRNA (Figure 7B), sequestering tumor-suppressive miRNAs including miR-10396b-5p, miR-486-3p, and miR-450a-2-3p. This miRNA sponging activity leads to reduced degradation of oncogenic mRNA targets, enhancing their expression. Together, these two mechanisms contribute to increased tumor cell proliferation and invasion, supporting a pro-tumorigenic role for TPL1 in TNBC.

**Figure 7.**
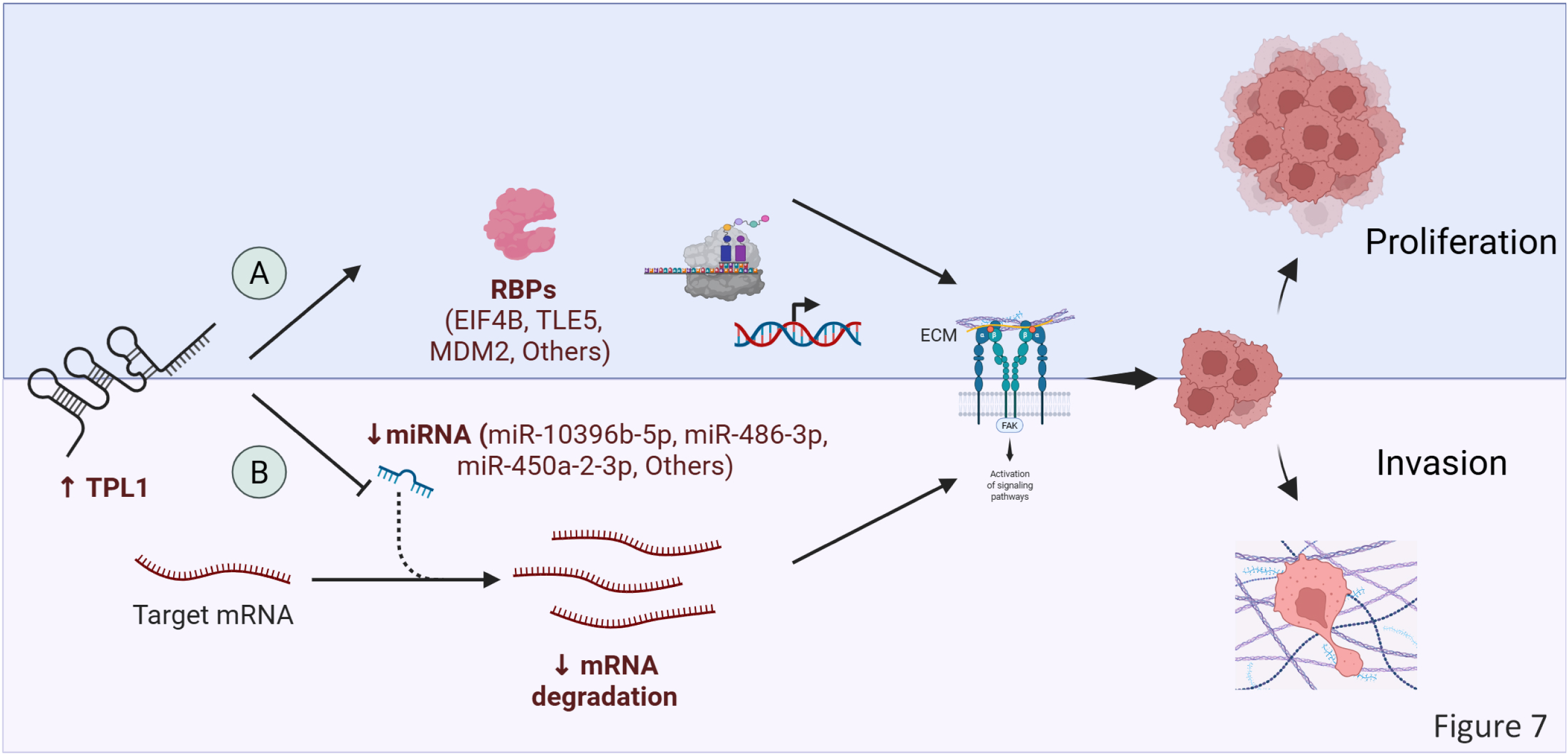
Schematic illustration of TPL1-mediated regulatory mechanisms in TNBC. **(A)** Upregulation of TPL1 promotes interaction with RNA-binding proteins (RBPs), including EIF4B, TLE5, and MDM2, potentially influencing transcriptional programs associated with proliferation and invasion. **(B)** TPL1 also acts as a competing endogenous RNA (ceRNA), sequestering specific miRNAs (e.g., miR-10396b-5p, miR-486-3p, miR-450a-2-3p), thereby preventing miRNA-mediated degradation of target mRNAs. This leads to stabilization of oncogenic transcripts and enhances invasive and proliferative phenotypes.

## Discussion

Advances in genomic research have uncovered a vast number of lncRNAs, surpassing the number of protein-coding genes (PCGs), yet the functions of most lncRNAs remain largely unknown. Our previous studies identified lncRNAs associated with breast cancer subtypes, including those promoting TNBC growth ^39,42–44^. However, investigating the function of individual lncRNAs is laborious and time-consuming, and thus a comprehensive catalog detailing the functional roles of most annotated lncRNAs is still lacking. High-throughput pooled CRISPR screens offer an unbiased approach to systematically explore the functions of both PCGs and lncRNAs across diverse biological processes. Our study advances this understanding by providing functional insights into lncRNAs involved in TNBC, highlighting their potential as therapeutic targets.

While previous studies primarily focused on annotated lncRNAs, our work distinguishes itself by systematically mapping the functional landscape of lncRNAs in TNBC, including both GENCODE-annotated and novel lncRNA candidates. We employed a robust CRISPR-del approach to identify lncRNAs essential for TNBC survival. This strategy enabled systematic functional interrogation of hundreds of lncRNAs, providing a comprehensive view of their roles in TNBC pathogenesis. CRISPR-del, known for enhanced perturbation efficiency and on-target specificity, proved a powerful tool, offering clear advantages over traditional methods ^45,46^. Targeting TSSs of lncRNA genes allowed efficient suppression while minimizing off-target effects. The CRISPR-del screen identified key lncRNAs influencing cell proliferation and sensitivity to doxorubicin, a commonly used chemotherapy for TNBC.

Among the novel candidates identified, TPL1 stood out due to its elevated expression in TNBC compared to normal breast tissue, with the highest levels observed in the aggressive BLIS TNBC subtype. Targeting TPL1 with ASOs demonstrated therapeutic benefits in both 2D and 3D culture models, showing minimal toxicity toward MCF10A immortalized breast epithelial cells (Figure S9), thus supporting the promise of RNA-based therapeutics. Interestingly, TPL1 suppression inhibited TNBC cell proliferation and invasion, as demonstrated using organ-on-chip high-content screening systems. Correlation analysis in TNBC tissue revealed strong associations between TPL1 and several hnRNP family members, suggesting a nuclear function for this lncRNA. To gain mechanistic insight into TPL1’s mode of action, transcriptomic and proteomic profiling revealed significant downregulation of pathways related to cell migration, cell adhesion, ECM-receptor interaction, PI3K-Akt signaling, and focal adhesion.

We further conducted a protein array to identify direct RBP interactors of TPL1, uncovering several significant partners. The cloned TPL1 transcript was 900 bp in length (Supplementary datafile1), starting from nucleotide 17 of the ENST00000747437.1 transcript, whereas the intended amplicon size was 1155 bp. Interestingly, this start site corresponded with the transcription start site identified in our merged assembly (MSTRG.37012.1). Notably, this gene has multiple transcript variants (Figure S10), all sharing the same first exon. The size of our cloned transcript may represent an alternative isoform that is currently not annotated in GENCODE, although it still contain the first common exon. Among the highest-ranking hits was EIF4B. Consistent with our findings, the gastric cancer metastasis-associated lncRNA GMAN has been shown to directly bind EIF4B and promote its phosphorylation, thereby stabilizing phosphorylated-EIF4B ^47^, contributing to increased proliferation and metastasis, mechanisms aligned with the phenotypic effects observed following TPL1 suppression in TNBC models. Additional TPL1 binding partners included TLE5, a transcriptional corepressor, and TARBP2, a double-stranded RNA-binding protein involved in miRNA processing. These interactions suggest TPL1 regulates gene expression at both transcriptional and post-transcriptional levels, contributing to its oncogenic functions in TNBC.

Other notable interactors included NACC1, a known oncogene in breast and ovarian cancer. Du et al. reported that LINC00319 accelerates ovarian cancer progression via the miR-423-5p/NACC1 axis ^48^, suggesting that TPL1 might exert similar effects through direct interaction with NACC1. MDM2, a key regulator of p53, was also found to interact directly with TPL1. Concordantly, Guo et al. reported that the non-coding repressor of NFAT (NRON) binds MDM2 to promote breast cancer tumorigenesis ^49^. Additional binding partners for TPL1 included MAP2K6, a component of the MAPK signaling cascade, indicating TPL1’s potential involvement in stress and growth signaling pathways. Other interactors of interest were RAB4A and MTMR2, involved in vesicle trafficking and phosphoinositide signaling, and DEPDC1 and PLEK2, which have roles in cell proliferation and migration. The identification of multiple RNA-binding proteins among the TPL1 interactors—such as HNRNPH1, NSUN4, PABPC1L2A, PABPC3, IGF2BP2, SRP19, MTRES1, NPM1, and RPL12—strongly supports a role for TPL1 in post-transcriptional gene regulation. These interactions suggest that TPL1 may function as a scaffold or regulator within RNA-protein complexes involved in splicing, mRNA stability, or translation. Notably, a subset of both high-confidence (Tier 1) and moderate-confidence (Tier 2) interactors—EIF4B, PFKM, STIP1, RAN, NOLC1, PRDX2, METTL2B, RPL14, NPM1, RPL12, and CACYBP—also showed positive correlation with TPL1 expression in TNBC patient datasets, suggesting transcriptional co-regulation or functional interdependence in vivo. The identification of RAN, a GTP-binding nuclear protein, further aligns with transcriptomic pathway data showing suppression of GTPase signaling upon TPL1 silencing, linking this lncRNA to intracellular trafficking and nuclear transport pathways. Our data are consistent with Cui et al. who reported lncRNA WFDC21P to directly bind RAN and to promote the Akt/GSK3β/β-catenin pathway activity in gastric cancer ^50^. Together, these findings position TPL1 at the intersection of RNA metabolism and oncogenic signaling, reinforcing its potential role as a functional hub in TNBC pathogenesis.

In parallel, we applied an integrative strategy to explore TPL1’s potential role as ceRNA by identifying miRNAs sponged by TPL1 and mapping their downstream mRNA targets. These findings highlight the potential role of specific miRNAs, such as miR-10396b-5p, miR-1285-3p, miR-1293, miR-23b-5p, miR-450a-2-3p, and miR-486-3p, in orchestrating critical cellular functions in TNBC. Their ability to regulate multiple transcripts involved in signal transduction (e.g., IQSEC2, HIPK2), cell adhesion and migration (e.g., ITGB4, CDCP1, LIMS2), and transcriptional control (e.g., TGFB1, TGFBR2, CIITA) suggests broad regulatory influence. The observation that several genes (SPRN, MDGA1, LHPP, CALB2) are targeted by multiple miRNAs further supports the presence of coordinated post-transcriptional control mechanisms, potentially reinforcing the robustness and specificity of gene regulation in TPL1-associated networks.

In summary, our study highlights the prognostic and therapeutic potential of lncRNAs in TNBC. By elucidating the functional roles of specific lncRNAs and their association with distinct TNBC subtypes, we have laid the foundation for further mechanistic studies and the development of novel, subtype-tailored therapeutic strategies. Future efforts to deepen our understanding of lncRNA-mediated regulatory networks in TNBC may enable identification of additional biomarkers and the creation of more effective targeted therapies, ultimately improving outcomes for patients with this aggressive breast cancer subtype.

## Limitations of the study

The CRISPR-del approach has inherent limitations, including the risk of false positives due to off-target effects. This risk is mitigated by employing approximately 10 pgRNAs per target and validating key findings using ASOs, which strengthens confidence in the screen’s reliability. Nonetheless, challenges remain. Targeting lncRNA’s TSSs may overlook certain lncRNAs, particularly those located near PCGs or those not amenable to TSS-directed targeting. To address these limitations, alternative CRISPR systems such as Cas13d/CasRx offer promising solutions by enabling RNA-targeted knockdown of lncRNAs ^51,52^. Some lncRNAs could be overlooked due to library design limitations or lower efficiency of CRISPR-del compared to other methods such as ORF mutations. Furthermore, our use of 2D monolayer cell models may not fully replicate the complex tumor microenvironment, highlighting the need for future studies incorporating 3D cultures and in vivo models. Moving forward, research should focus on clinically relevant lncRNAs, particularly those associated with residual disease and poor prognosis, as these are promising candidates for RNA-based therapeutic development.

## Supporting information

Supplemental datafile 1

## Acknowledgment

This work was supported by funds from Qatar National Research Fund (grant no. NPRP12S-0221-190124) and from Qatar Biomedical Research institute (QBRI) intramural grant program (IGP5-2022-006) for Dr. Nehad M. Alajez. This work was partially supported by Science Foundation Ireland through Future Research Leaders award 18/FRL/6194 (RJ). M.C. acknowledges the support of DevelopMed, which received funding from the European Union’s Horizon 2020 research and innovation programme under the Marie Skłodowska-Curie grant agreement No 945425. This research was funded by Science Foundation Ireland under Grant number [18/CRT/6214] (SR) and in part by the EU’s Horizon 2020 research and innovation programme under the Marie Sklodowska-Curie grant H2020-MSCA-COFUND-2019-945385 (MC). YY acknowledges the financial support from China Scholarship Council program. We thank Dr. Houari B. Abdesselem and Ms. Israa E Elbashir (HBKU Proteomic Core Facility, Hamad Bin Khalifa University) for the protein microarray experiments.

## Declarations

### Ethical Approval

Not applicable

### Authors’ contributions

R.E., performed the experiments and wrote the manuscript; S.R., designed the pgRNA library; R.V., L.Y.O., C.S., F.S., and A.A.H.Z, conducted experiments and data analysis; M.C. and T.U., assisted with experiments; Y.Y., performed bioinformatics analysis and hit identification; C.P.Q., constructed the merged assembly; K.O., carried out genomic PCR and NGS; R.J. contributed to the concept and design of the study; N.A., conceptualized the study, secured funding, and approved the final manuscript.

### Availability of data and materials

All processed data are provided in supplementary data. The accession numbers of datasets analyzed in the current study are indicated in the methods section. Additional data is available upon request from the corresponding author.

