## Supplemental Table 2 for "Robust CRISPR Screens Identify TPL1 as a Novel Long Noncoding RNA Driving Triple-Negative Breast Cancer Hallmarks"

| **Table S2. Primer List for Pathway Validation, Gene Knockdown, Full Length and Genotyping.** | | | |
| --- | --- | --- | --- |
| **No** | Target name | **Forward Sequences** | **Reverse Sequences** |
| 1 | β-ACTIN | 5′-GGCACCCAGCACAATGAAG-3′ | 5′-CCGATCCACACGGAGTACTTG-3′ |
| 2 | ITGA3 | 5′-TGTGGCTTGGAGTGACTGTG-3′ | 5′-TCATTGCCTCGCACGTAGC-3′ |
| 3 | ITGB8 | 5′-GTGAAAGTCATATCGGATGGCG-3′ | 5′-GCTATCAAGAGCGAGATGAGACG-3′ |
| 4 | ITGB4 | 5′-CTCCACCGAGTCAGCCTTC-3′ | 5′-CGGGTAGTCCTGTGTCCTGTA-3′ |
| 5 | ITGB5 | 5′-AACTCGCGGAGGAGATGAG-3′ | 5′-GGTGCCGTGTAGGAGAAAGG-3′ |
| 6 | COL6A3 | 5′-ATGAGGAAACATCGGCACTTG-3′ | 5′-GGGCATGAGTTGTAGGAAAGC-3′ |
| 7 | ITGA10 | 5′-ACTTAGGTGACTACCAACTGGG-3′ | 5′-CCACAAGCACGAGACCAGA-3′ |
| 8 | COL4A5 | 5′-CAAAAGGTGATCGTGGTTTCCC -3′ | 5′-GTCCAGGTTGTCCATTTGGTC-3′ |
| 9 | LAMA3 | 5′-CACCGGGATATTTCGGGAATC -3′ | 5′-AGCTGTCGCAATCATCACATT-3′ |
| 10 | PRKCA | 5′-GTCCACAAGAGGTGCCATGAA -3′ | 5′-AAGGTGGGGCTTCCGTAAGT-3′ |
| 11 | RPTOR | 5′-AATGCTGCAATCGCCTCTTCT -3′ | 5′-GCCAAAGGTAGGTTCCAGTCTG-3′ |
| 12 | TLN2 | 5′-GCGTGTCGAGTCATTCGGG -3′ | 5′-CCCTTTCCTCGGGTCTTCATC-3′ |
| 13 | PTK7 | 5′-GTAGTAGCGAGGTATGAGGAGG -3′ | 5′-TGCGGTTAGTGATGGGAGTCT-3′ |
| 14 | CAPN2 | 5′-GTTCTGGCAATACGGCGAGT -3′ | 5′-CTTCGGCTGAATGCACAAAGA-3′ |
| 15 | TGFB1 | 5′-CAATTCCTGGCGATACCTCAG-3′ | 5′-GCACAACTCCGGTGACATCAA-3′ |
| 16 | DOCK1 | 5′-ACCGAGGTTACACGTTACGAA-3′ | 5′-TCGGAGTGTCGTGGTGACTT-3′ |
| 17 | RBM6 | 5′-TGGAGTATGTATCAAGCCTGGA-3′ | 5′-ATGAACAGGAAGATCGGTGCC-3′ |
| 18 | AGO2 | 5′-CCATGTACTCGGGAGCCG-3′ | 5′-TCCCAAAGTCGGGTCTAGGT-3′ |
| 19 | VEGFA | 5′-CTGTCTAATGCCCTGGAGCC-3′ | 5′-GTCACATCTGCAAGTACGTTCG-3′ |
| 20 | MMP14 | 5′-CGAGGTGCCCTATGCCTAC-3′ | 5′-CTCGGCAGAGTCAAAGTGG-3′ |
| 21 | PLAU | 5′-GGGAATGGTCACTTTTACCGAG-3′ | 5′-GGGCATGGTACGTTTGCTG-3′ |
| 22 | THBS2 | 5′-GACACGCTGGATCTCACCTAC-3′ | 5′-GAAGCTGTCTATGAGGTCGCA-3′ |
| 23 | FN1 | 5′-CGGTGGCTGTCAGTCAAAG-3′ | 5′-AAACCTCGGCTTCCTCCATAA-3′ |
| 24 | Cas9 | 5′-CTGGACGCCACCCTGATCC-3′ | 5′-GACGCTGTAGTCTTCAGAGATG-3′ |
| 25 | TPL1 | 5′-TCTTCTACACGCCACTTCGC-3′ | 5′-TCCATCTTTCATTCTCCTTTGATGT-3′ |
| 26 | PROLIT1 | 5′-TCAGAGTCCACAGGATGGATG-3′ | 5′-GGGATGTTCAGGCTTGATCAGAAA-3′ |
| 27 | TPL1 Full Length | 5′-TAATACGACTCACTATAGGGCAGCTAAG AGGGTGCGAGAT-3′ | 5′-CAAGGCAGAAGAATTTTTCTTAGTATA GAAC-3′ |
| 28 | Candidate 256 (LINC00910) | 5′-CCACTGGTGACGACGTAAAG-3′ | 5′-GCCAAGATTGCGCTACTACA-3′ |
| 29 | Candidate 256_genotyping | 5′-ACTACCTTGCCTAGGCTGTTTC-3′ | 5′-CGTGTGTCGTGCTCATTACG-3′ |
