## Supplemental Figure 1 for "Robust CRISPR Screens Identify TPL1 as a Novel Long Noncoding RNA Driving Triple-Negative Breast Cancer Hallmarks"

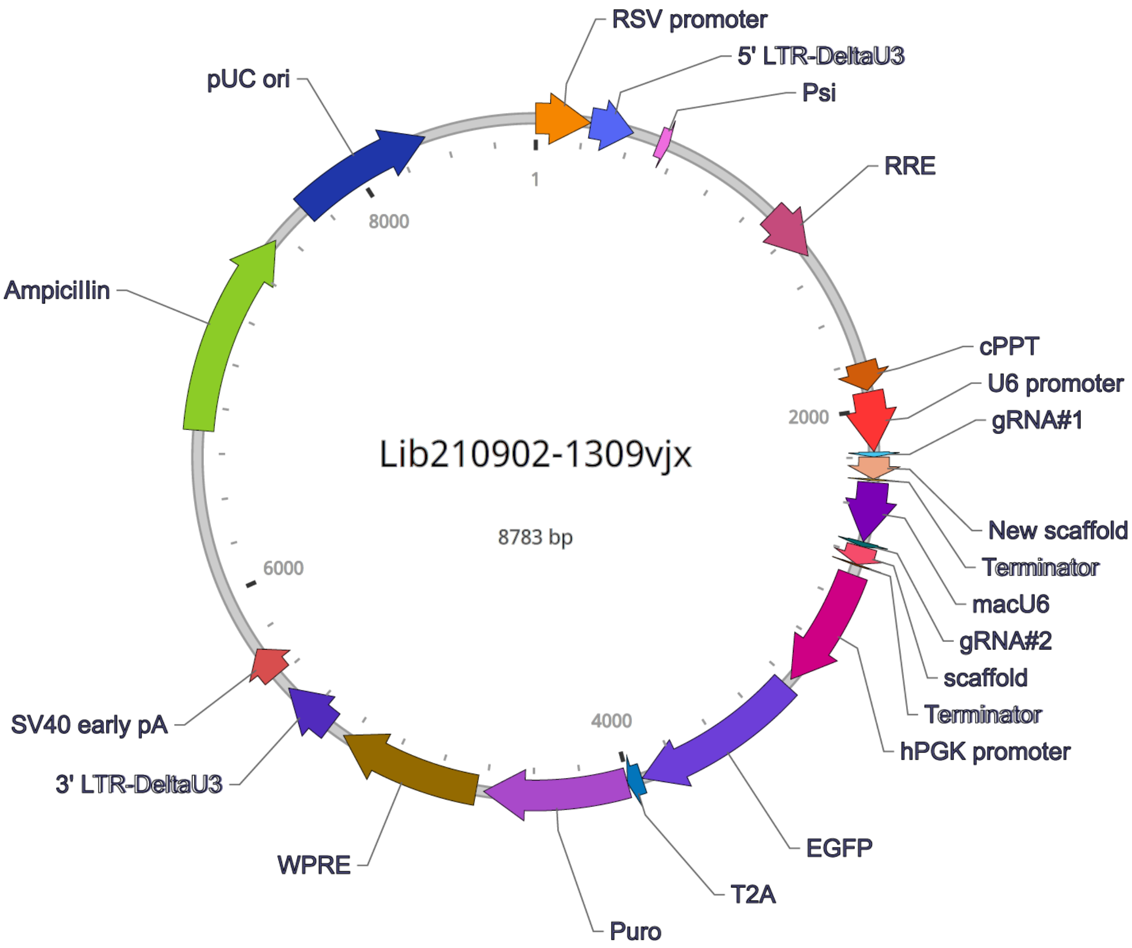


**Figure S1. Map depicting key features of the mammalian pgRNA expression lentiviral vector used for library cloning.**
