## Supplementary figures and images for "Robust CRISPR Screens Identify TPL1 as a Novel Long Noncoding RNA Driving Triple-Negative Breast Cancer Hallmarks"

### Supplemental Figure 2

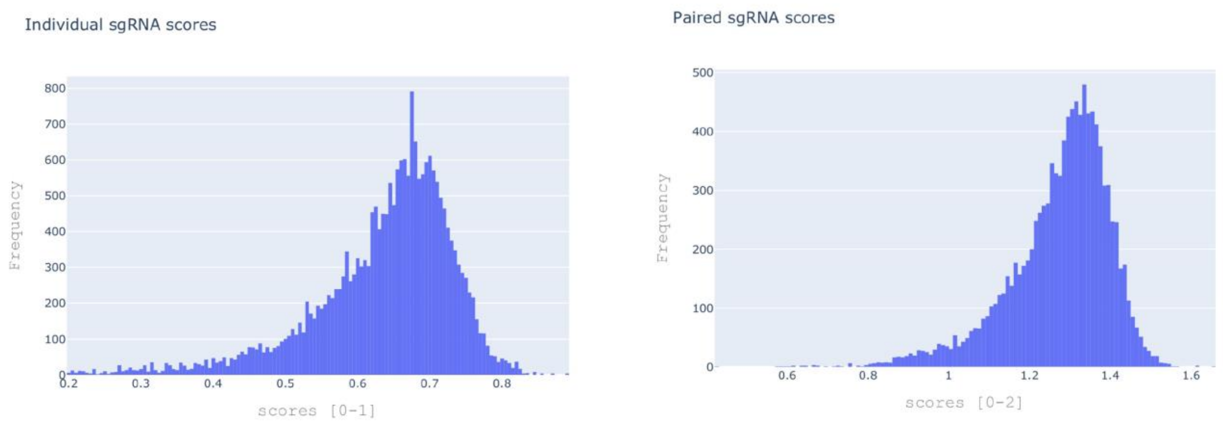


**Figure S2. Individual and pgRNA scores from the pgRNA library.**
