## Supplemental Figure 3 for "Robust CRISPR Screens Identify TPL1 as a Novel Long Noncoding RNA Driving Triple-Negative Breast Cancer Hallmarks"

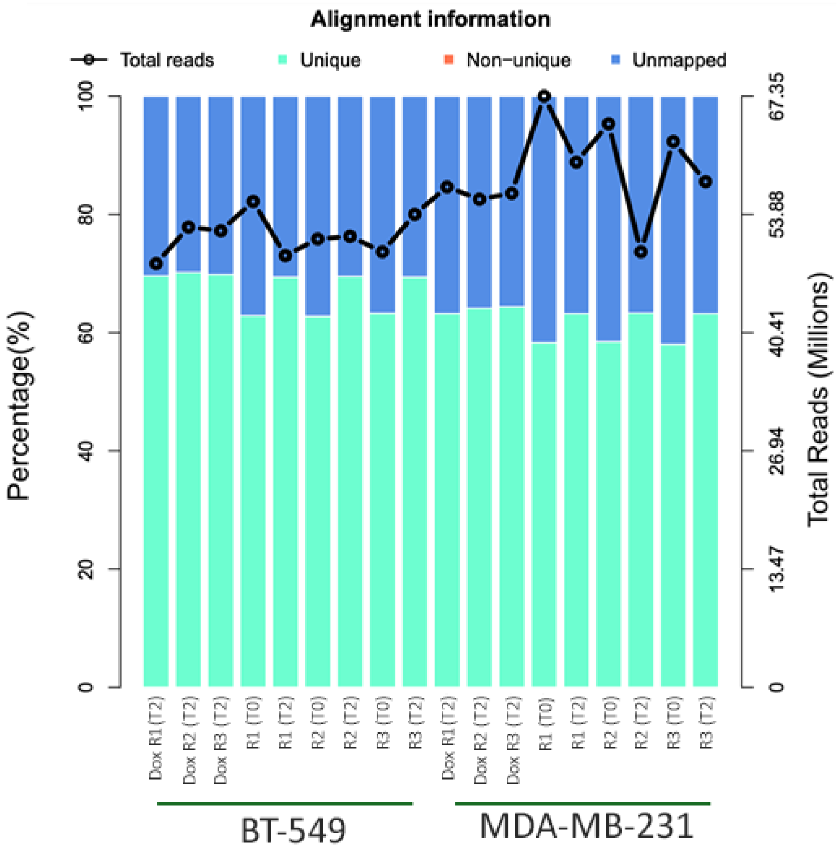


**Figure S3. Percent alignment of NGS genomic amplicons to pgRNA library sequences from MDA-MB-231 and BT-549 TNBC cells.**
