## Supplemental Figure 4 for "Robust CRISPR Screens Identify TPL1 as a Novel Long Noncoding RNA Driving Triple-Negative Breast Cancer Hallmarks"

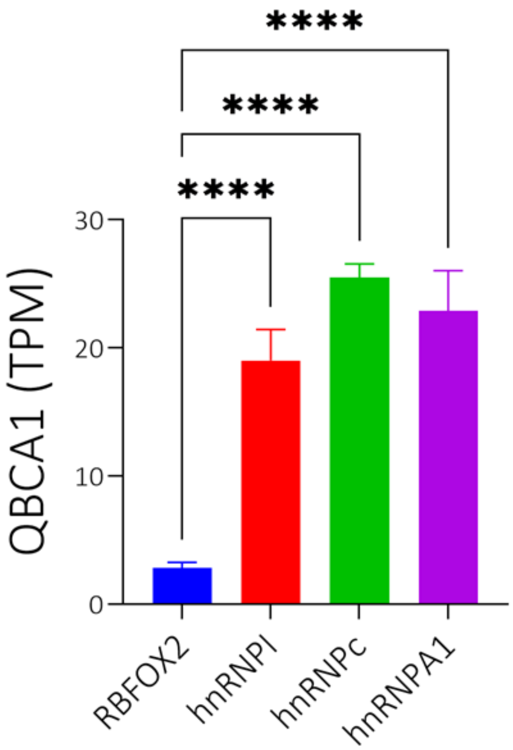


**Figure S4. TPL1 is enriched in hnRNP-bound RNA fractions compared to RBFOX2. RNA-Seq data for** RNA immunoprecipitation (hnRNPA1, hnRNPAI and hnRNPC, and RBFOX2) followed by sequencing was retrieved from PRJNA935334 dataset. RNA-Seq were then mapped to GENCODE R 47 and the association (transcript per million, TPM) was estimated. Strong enrichment of QBCA1 was observed in hnRNPI, hnRNPA1 and hnRNPC, while RBFOX2-IP showed minimal signal. Data re presented as mean ± S.E. **** $p < 0.00005$.
