## Supplemental Figure 5 for "Robust CRISPR Screens Identify TPL1 as a Novel Long Noncoding RNA Driving Triple-Negative Breast Cancer Hallmarks"

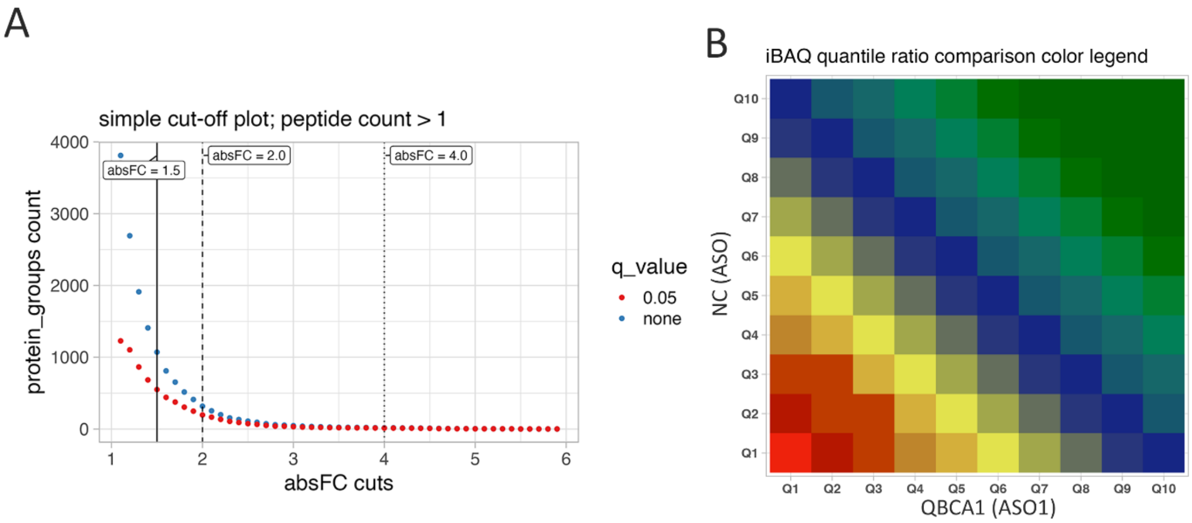


**Figure S5. Quantitative proteomic profiling of TPL1-depleted TNBC cells. (A)** Simple cut-off plot illustrating the distribution of differentially expressed proteins based on absolute fold change (x-axis) and the number of protein groups (y-axis). Vertical lines correspond to absolute fold changes (absFC) of 1.5, 2.0, and 4.0. The red line indicates a q-value of 0.05, and the blue line indicates no q-value threshold. **(B)** iBAQ quantile ratio comparison between TNBC cells transfected with negative control (NC) ASO (x-axis) and TPL1-targeting ASO (y-axis), based on DIA-MS data. Each cell represents a matched quantile-to-quantile comparison (Q1–Q10) of protein abundance. Warmer colors (red/yellow) indicate proteins more abundant in NC ASO-treated cells, while cooler colors (blue/green) indicate proteins enriched in TPL1-depleted cells. n = 5 biological replicates per group.
