## Supplemental Figure 6 for "Robust CRISPR Screens Identify TPL1 as a Novel Long Noncoding RNA Driving Triple-Negative Breast Cancer Hallmarks"

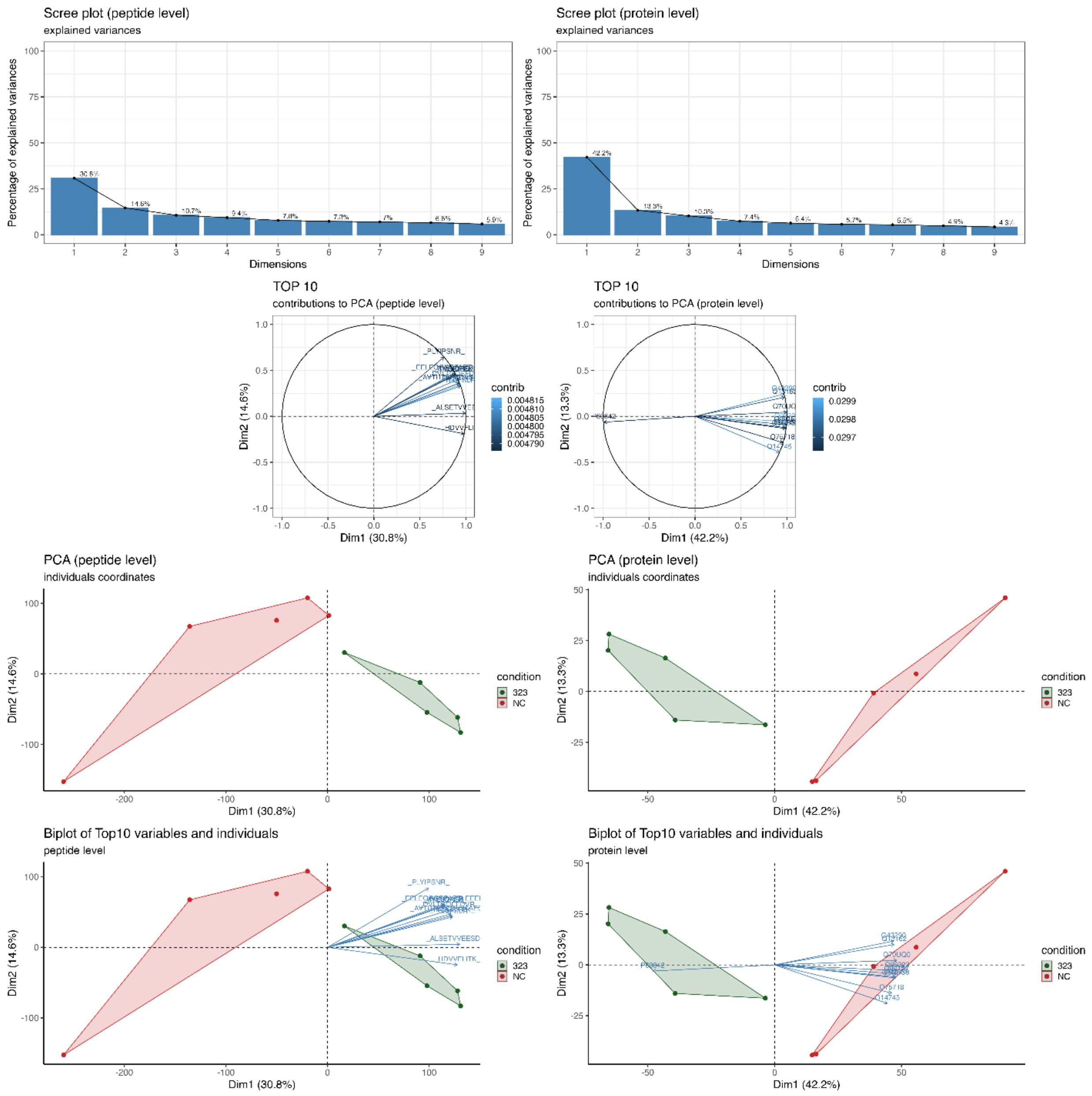


**Figure S6. Principal Component Analysis (PCA) of TPL1 protein array data at peptide and protein levels.** PCA was performed on normalized intensity values from the human proteome array following hybridization with in vitro transcribed TPL1 RNA. (Top) Scree plots showing the percentage of variance explained by each principal component at the peptide (left) and protein (right) levels. (Middle) Contribution plots displaying the top 10 most influential variables contributing to PC1 and PC2 at both levels. (Bottom) PCA biplots show the distribution of experimental conditions (323: TPL1, NC: negative control) along the first two principal components, highlighting sample separation and variable contributions. Distinct clustering of conditions reflects differential TPL1–protein interactions.
