## Supplemental Figure 7 for "Robust CRISPR Screens Identify TPL1 as a Novel Long Noncoding RNA Driving Triple-Negative Breast Cancer Hallmarks"

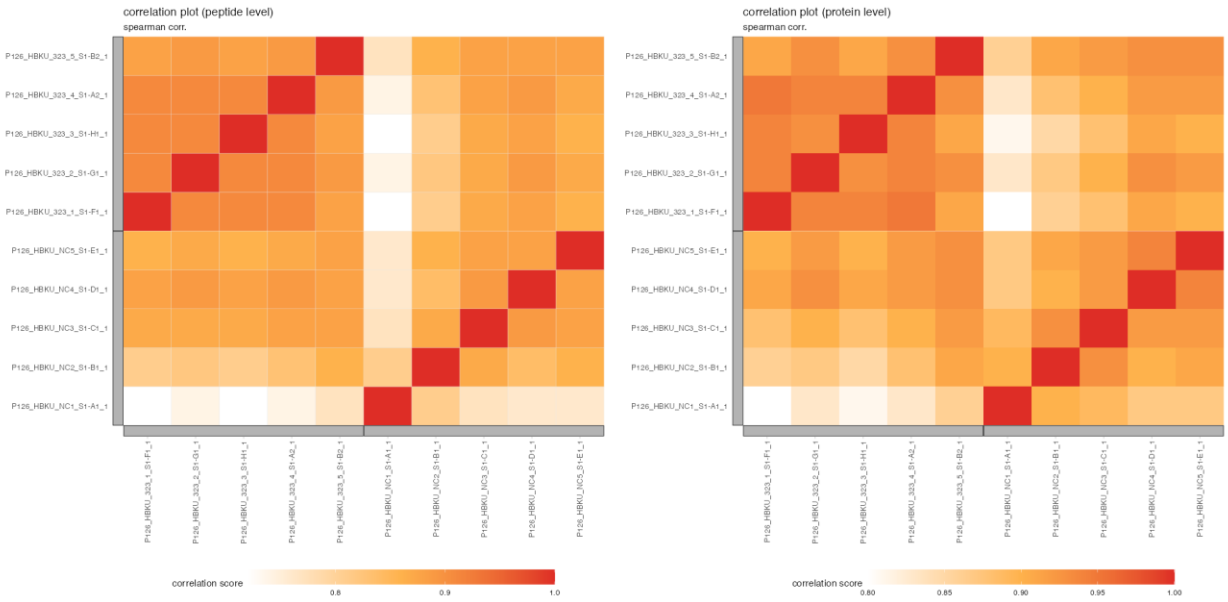


**Figure S7. Correlation heatmaps of proteome array samples at peptide and protein levels.** Spearman correlation plots showing pairwise relationships between biological replicates from TPL1-transfected (323) and negative control (NC) conditions. Heatmaps were generated at both the peptide level (left) and protein level (right), demonstrating strong intra-group correlations and clear separation between experimental conditions. Higher correlation among replicates within each group supports the reproducibility of the protein array experiment and distinct binding profiles associated with TPL1.
