## Supplemental Figure 8 for "Robust CRISPR Screens Identify TPL1 as a Novel Long Noncoding RNA Driving Triple-Negative Breast Cancer Hallmarks"

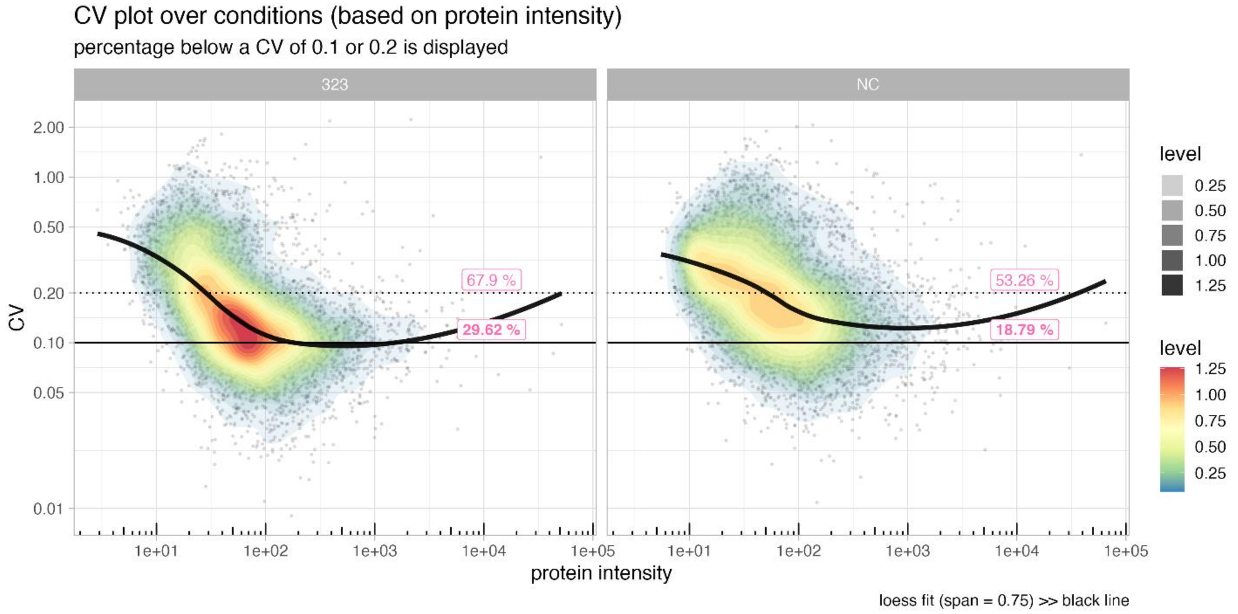


**Figure S8. Coefficient of variation (CV) versus protein intensity for TPL1 (323) and negative control (NC) conditions.** CV plots display the distribution of protein-level variability (CV) as a function of signal intensity for each condition. Density color maps represent local data point density, and the black line indicates a LOESS fit (span = 0.75). Dashed and solid horizontal lines represent CV thresholds of 0.2 and 0.1, respectively.
