## Supplemental Figure 9 for "Robust CRISPR Screens Identify TPL1 as a Novel Long Noncoding RNA Driving Triple-Negative Breast Cancer Hallmarks"

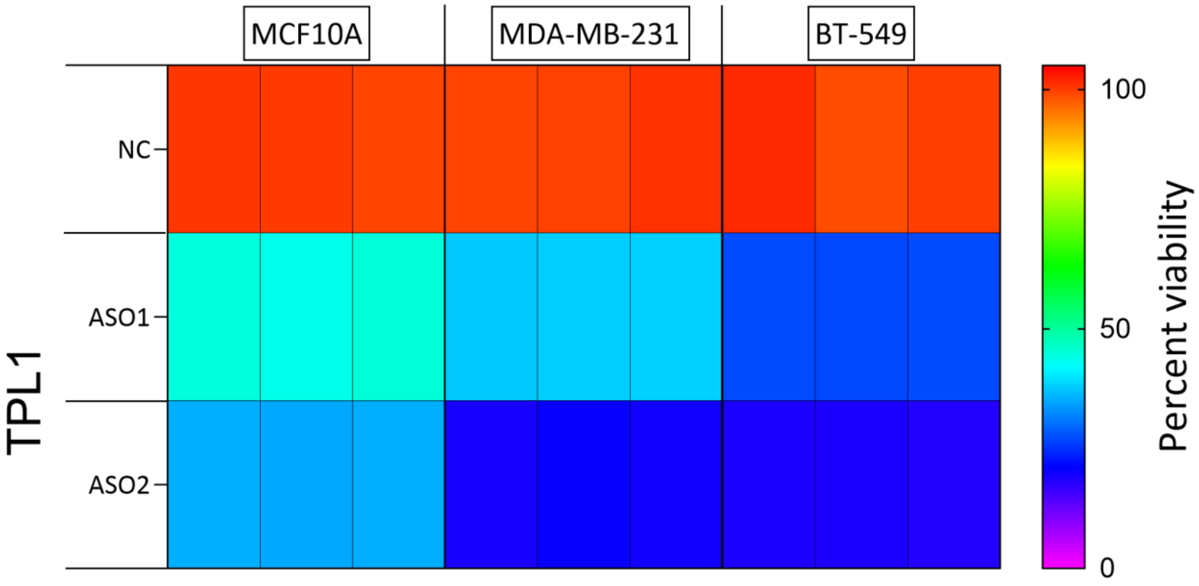


**Figure S9. Summary of the effects of TPL1-targeting ASOs and negative control ASO on TNBC and normal breast cell lines.** The figure presents a heatmap summarizing the impact of antisense oligonucleotides (ASOs) targeting TPL1 on cell viability across various triple-negative breast cancer (TNBC) and normal breast epithelial cell lines. Rows represent the lncRNA target (TPL1), while columns indicate different ASO designs and cell lines. Cell viability is expressed as a percentage normalized to the negative control (NC) ASO treatment in each respective cell line.
