## Supplemental Figure 10 for "Robust CRISPR Screens Identify TPL1 as a Novel Long Noncoding RNA Driving Triple-Negative Breast Cancer Hallmarks"

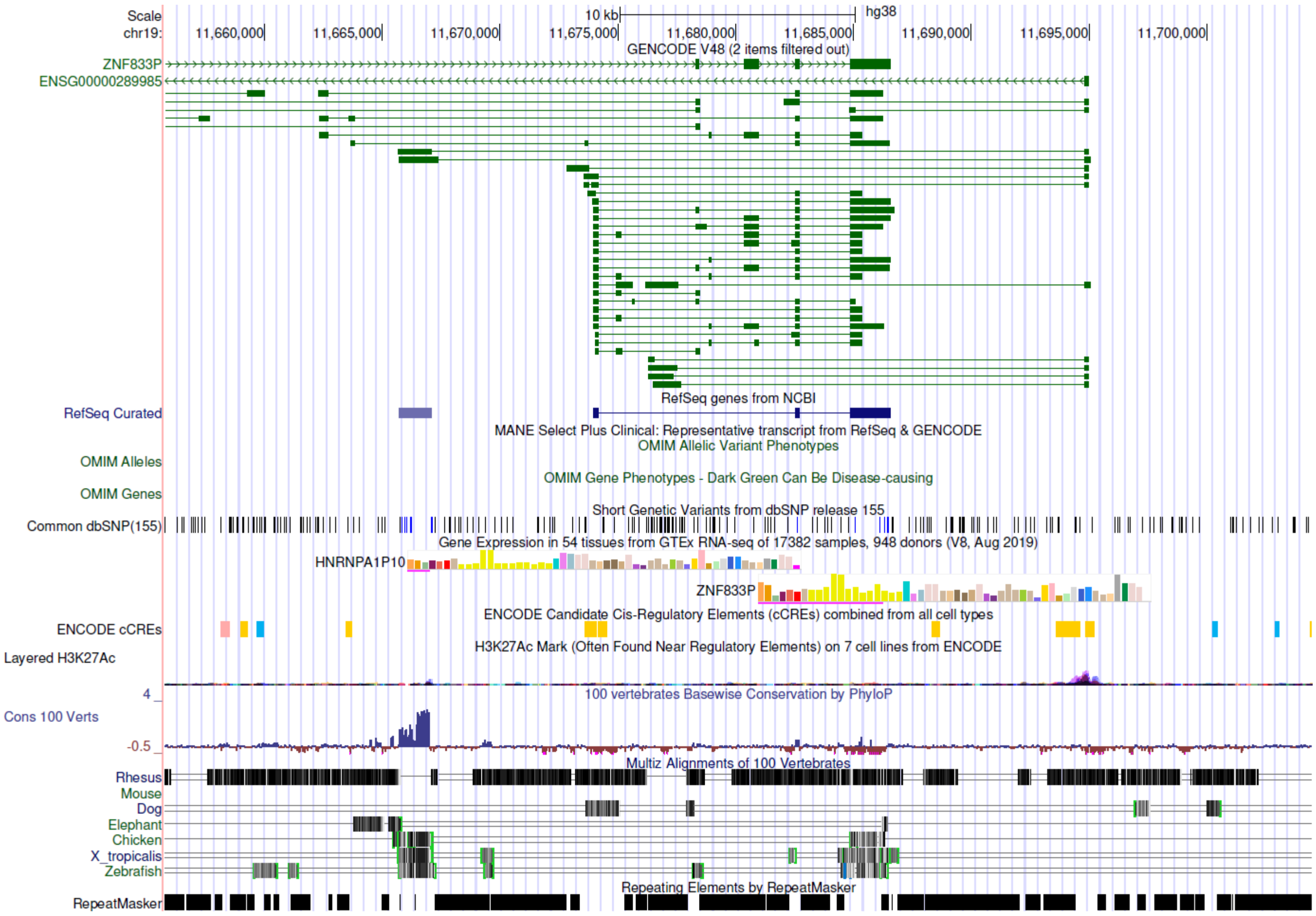


**Figure S10. Genomic coordinates and transcript isoforms of TPL1 (ENSG00000289985) visualized using the UCSC Genome Browser.** The figure displays the genomic locus of the lncRNA TPL1 (ENSG00000289985) on the human reference genome (hg38), as visualized in the UCSC Genome Browser. All annotated transcript isoforms are shown, including exon–intron structures, demonstrating the alternative splicing complexity of the TPL1 locus.
