## Supplemental datafile 1 for "Robust CRISPR Screens Identify TPL1 as a Novel Long Noncoding RNA Driving Triple-Negative Breast Cancer Hallmarks"

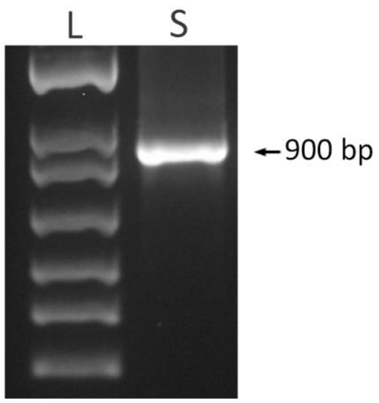


**RT-PCR amplification of TPL1 (ENST00000702307).**

**TPL1 Sanger sequence**

CAGCTAAGAGGGTGCGAGATTGAAAAAGCCCGCGGCGTCTTCTACACGCCACTTCGCTGGGCTCCTGTCT
CTGTCGCTGAGAGGCACTCTGGAGAGTCTGCGGCAGCCTCTGTCACACAGGGACCTGCACTGGTCGTGGG
AGCCGCAGAGAGGCCCCCGGGACACGGAGGAATTCGGAAAATGGCTGGAGTGCAGTGGCGTGACCTGGGC
TTGCTACAACCTCCACCTCCCAGCCGCCTGCCTTGGCCTCCCAAAGTGCCGAGATTGCAGCCTCTGCCCG
GCCGCCACCCCGTCTGGGAAGTGAGGAGCGTCTCTGCCTGGCCGCCCATCATCTGGGATGCGAGGAGCCC
CTCTGCCCGGCTGCCCAGTCTGGGAAGTGAGGAGCGCCTCTTCCCGGCCACCATCCCATCTAGGAAGTGA
GGAGTGTCTCTGCCTGGCCGCCCATGGTCTGGGATGTGAGGAGCGCCTCTGCCCTGCTGCCCAGTCTGGG
AAGTGAGGAGCGCCTCCTCCCGGCGCCATCCCATCTAGGAAGTGAGGAGCGTCTCTGCCTTGCCGCCCAT
CGTCTGAGATGTGGGGAGCGCCTCTGCCCCACTGCCCTGTCTAGGATGTGAGGAGCGCCTCTACCCGGCC
GTGACCCCGTCTGGGAACTGAGGAGTGTCTCTGCCTGACCACCACCCCGTCTGGGAGGTGACGAGCGCCT
CCGCCTGGCCTCCACCCCGTCTGGGAGGTGTACCCAACAGCTCACTGAGAACGGGCCATGATGACGATGG
CAGTTTTGTCGAATAGAAAAGGGGGAAATGTGGGGAAAAGAAAGATAAGATTGTTACTGTGTCTGTGTAG
ATAGTAGTAGACATAGGAGACTCCATTTTGTTCTATACTAAGAAAAATTCTTCTGCCTTG
